# Map of spiking activity underlying change detection in the mouse visual system

**DOI:** 10.1101/2025.10.17.683190

**Authors:** Corbett Bennett, Sam Gale, Gregg Heller, Tamina Ramirez, Hannah Belski, Alex Piet, Omid Zobeiri, Adam Amster, Anton Arkhipov, Alex Cahoon, Shiella Caldejon, Mikayla Carlson, Linzy Casal, Scott Daniel, Colin Farrell, Marina Garrett, Ryan Gillis, Conor Grasso, Ben Hardcastle, Ross Hytnen, Tye Johnson, Peter Ledochowitsch, Quinn L’Heureux, Dana Mastrovito, Ethan McBride, Stefan Mihalas, Chris Mochizuki, Chris Morrison, Chelsea Nayan, Kiet Ngo, Kat North, Doug Ollerenshaw, Ben Ouellette, Paul Rhoads, Kara Ronellenfitch, Martin Schroedter, Joshua H. Siegle, Cliff Slaughterbeck, David Sullivan, Jackie Swapp, Mike Taormina, Wayne Wakeman, Xana Waughman, Ali Williford, John Phillips, Peter A. Groblewski, Severine Durand, Christof Koch, Shawn R. Olsen

## Abstract

Visual behavior requires coordinated activity across hierarchically organized brain circuits. Understanding this complexity demands datasets that are both large-scale (sampling many areas) and dense (recording many neurons in each area). Here we present a database of spiking activity across the mouse visual system—including thalamus, cortex, and midbrain—while mice perform an image change detection task. Using Neuropixels probes, we record from >75,000 high-quality units in 54 mice, mapping area-, cortical layer-, and cell type-specific coding of sensory and motor information. Modulation by task-engagement increased across the thalamocortical hierarchy but was strongest in the midbrain. Novel images modulated cortical (but not thalamic) responses through delayed recurrent activity. Population decoding and optogenetics identified a critical decision window for change detection and revealed that mice use an adaptation-based rather than image-comparison strategy. This comprehensive resource provides a valuable substrate for understanding sensorimotor computations in neural networks.

## Introduction

A central challenge in neuroscience is to understand how diverse neurons, connected through distributed circuits, transform sensory information into behavior. The mammalian visual system exemplifies this complexity—visual processing engages hierarchical interactions across many brain regions including thalamus, cortex, and midbrain, with each local circuit containing heterogeneous cell types that coordinate through intricate connectivity to guide behavior. These visual circuits must be both robust (maintaining learned sensorimotor associations) and flexible (adapting to experience as animals encounter novel environments). To understand how spiking activity in the visual system implements adaptable sensorimotor computations, we need comprehensive datasets that measure the activity of multiple brain regions and cell types simultaneously during visually guided behavior.

Recent technological advances, including Neuropixels probes, have revolutionized our ability to record from large populations of neurons simultaneously^1–6^. Indeed, several large-scale experimental efforts in mice have already begun to systematically map brain-wide neural activity during sensory processing and decision-making^7–10^. For the most part, these recordings have been made serially (with one or two probes at a time) and shallowly (sampling tens to hundreds of well-isolated neurons total from each area). While such surveys have provided crucial insights into distributed neural computation^7–10^, understanding how a particular network coordinates activity on the level of cell-types and cortical layers requires a complementary approach: dense, simultaneous recordings from highly interconnected regions.

At the Allen Institute, we previously built standardized physiological pipelines for dense sampling of the cortical and thalamic visual system^4,11–13^. Our prior Visual Coding Neuropixels dataset surveyed stimulus-evoked spiking activity in passively viewing mice, and through our own analysis^4,14,15^ and dozens of follow-up studies from other groups^16^, has yielded many insights into neural processing in the cortex, thalamus and hippocampus. However, this dataset lacks a behavioral task and is thus ill-suited to studying sensorimotor transformations.

Here we describe a large-scale open dataset in which we used multi-regional Neuropixels recordings to map spiking activity across the mouse visual system while mice performed a visual change detection task. Change detection is a fundamental visual computation essential for survival, often involving rapid decision-making following a salient change in the environment—the sudden appearance of a predator, a child chasing a ball into the street. Recent work in the mouse has begun to identify circuits in the visual system involved in this process^8,12,17–22^, but a comprehensive map of spiking activity across multiple levels of the visual thalamocortical hierarchy is lacking.

Our experimental design incorporated several key features: (1) simultaneous recordings from six visual cortical areas at retinotopically matched locations^1^, (2) dense sampling across cortical layers and identification of genetically-defined interneuron subclasses through optotagging, (3) recordings from visual thalamic and midbrain regions known to be involved in visual decision making, and (4) a multi-epoch experimental structure capturing neural activity during active behavior, passive stimulus replay, and processing of both novel and familiar stimuli.

This resource—the Visual Behavior Neuropixels dataset—is readily available on public cloud platforms (Figure S1, Table 1) together with comprehensive tutorials and documentation (Table 2), enabling investigation of fundamental questions about neural computation in the visual system. In this study, we establish a foundation for future inquiry by 1) characterizing the diversity of sensory and motor responses in individual units and how these responses are modulated by task-relevant stimuli and engagement, 2) establishing the critical time window during which mice make a change detection decision and elucidating the computational strategy they use to detect change, and 3) describing the area-, cortical layer- and cell type-specific circuits through which novelty modulation arises across the visual hierarchy.

**Table 1:**
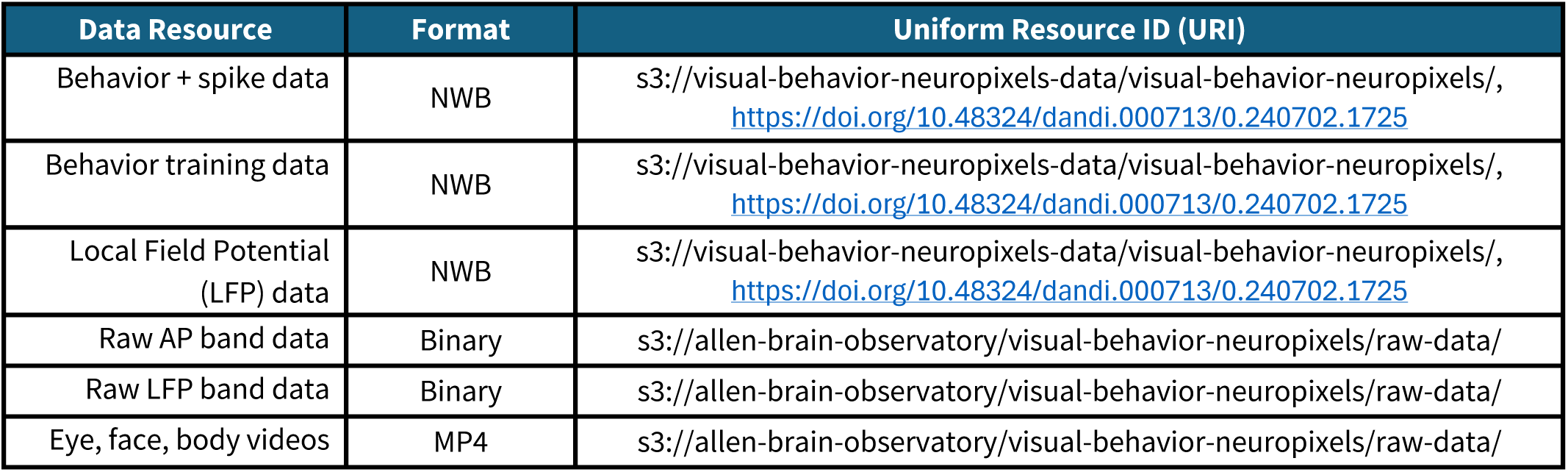
Data Resources associated with this dataset.

**Table 2:**
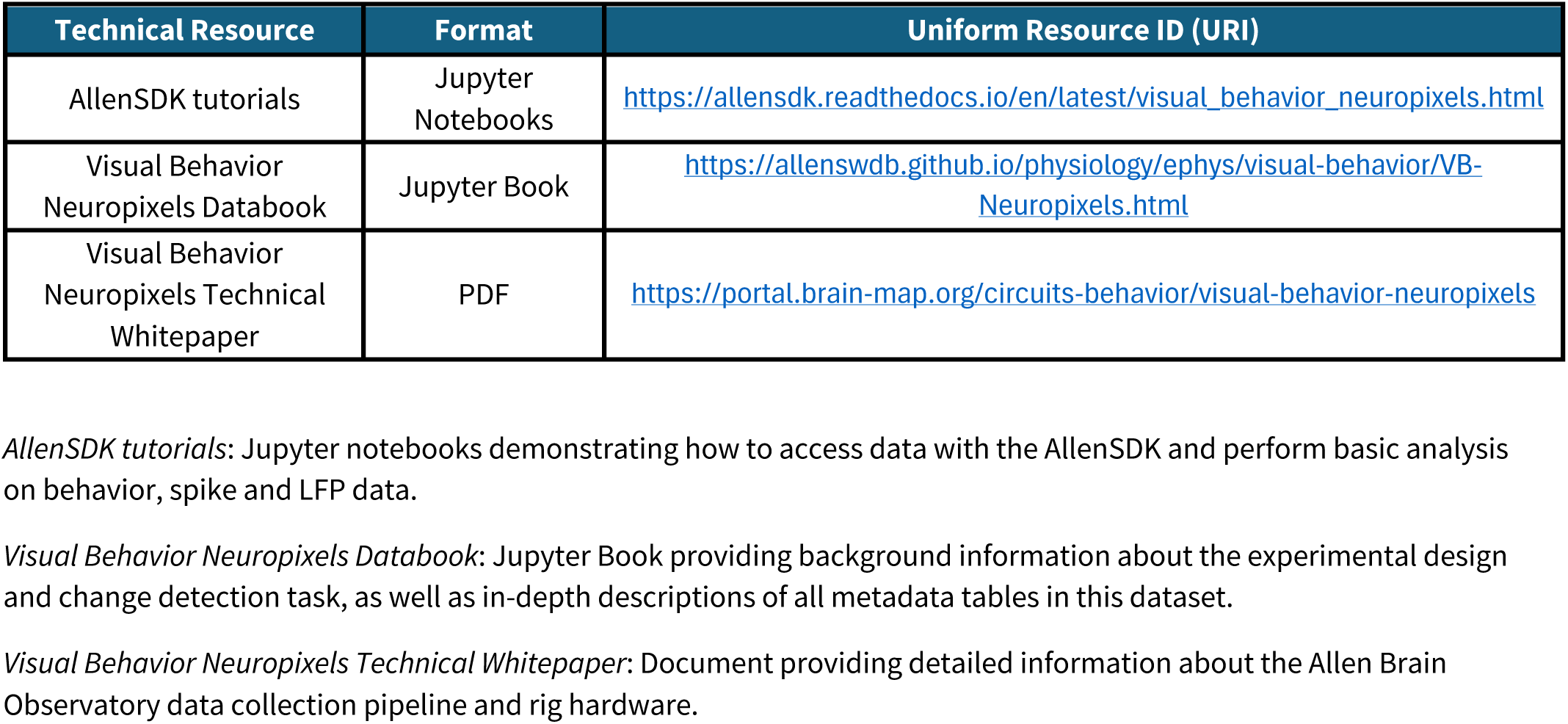
Technical Resources associated with this dataset.

| Technical Resource | Format | Uniform Resource ID (URI) |
| --- | --- | --- |
| AllenSDK tutorials | Jupyter Notebooks | https://allensdk.readthedocs.io/en/latest/visual_behavior_neuropixels.html |
| Visual Behavior Neuropixels Databook | Jupyter Book | https://allenswdb.github.io/physiology/ephys/visual-behavior/VB-Neuropixels.html |
| Visual Behavior Neuropixels Technical Whitepaper | PDF | https://portal.brain-map.org/circuits-behavior/visual-behavior-neuropixels |

*AllenSDK tutorials:* Jupyter notebooks demonstrating how to access data with the AllenSDK and perform basic analysis on behavior, spike and LFP data.
*Visual Behavior Neuropixels Databook:* Jupyter Book providing background information about the experimental design and change detection task, as well as in-depth descriptions of all metadata tables in this dataset.
*Visual Behavior Neuropixels Technical Whitepaper:* Document providing detailed information about the Allen Brain Observatory data collection pipeline and rig hardware.

## Results

### Survey of spiking activity during visual change detection

We recorded spiking data simultaneously from six Neuropixels probes, each targeting a distinct visual cortical region (Figure 1a; targets: VISp, VISl, VISal, VISrl, VISam, VISpm)^1,4^. Before the experiment, intrinsic signal imaging was used to map the borders and retinotopy of each cortical area; these maps then guided probe insertions to record from neurons with receptive fields near the center of the stimulus monitor (Figure 1a,b). Probes were inserted 3-3.5 mm into the brain to record subcortical regions (including thalamus, midbrain, and hippocampus) in addition to visual cortex (Figure 1c,d). In total, this dataset includes 76,091 high-quality units from 54 mice over 103 behavioral sessions (see Methods).

**Figure 1:**
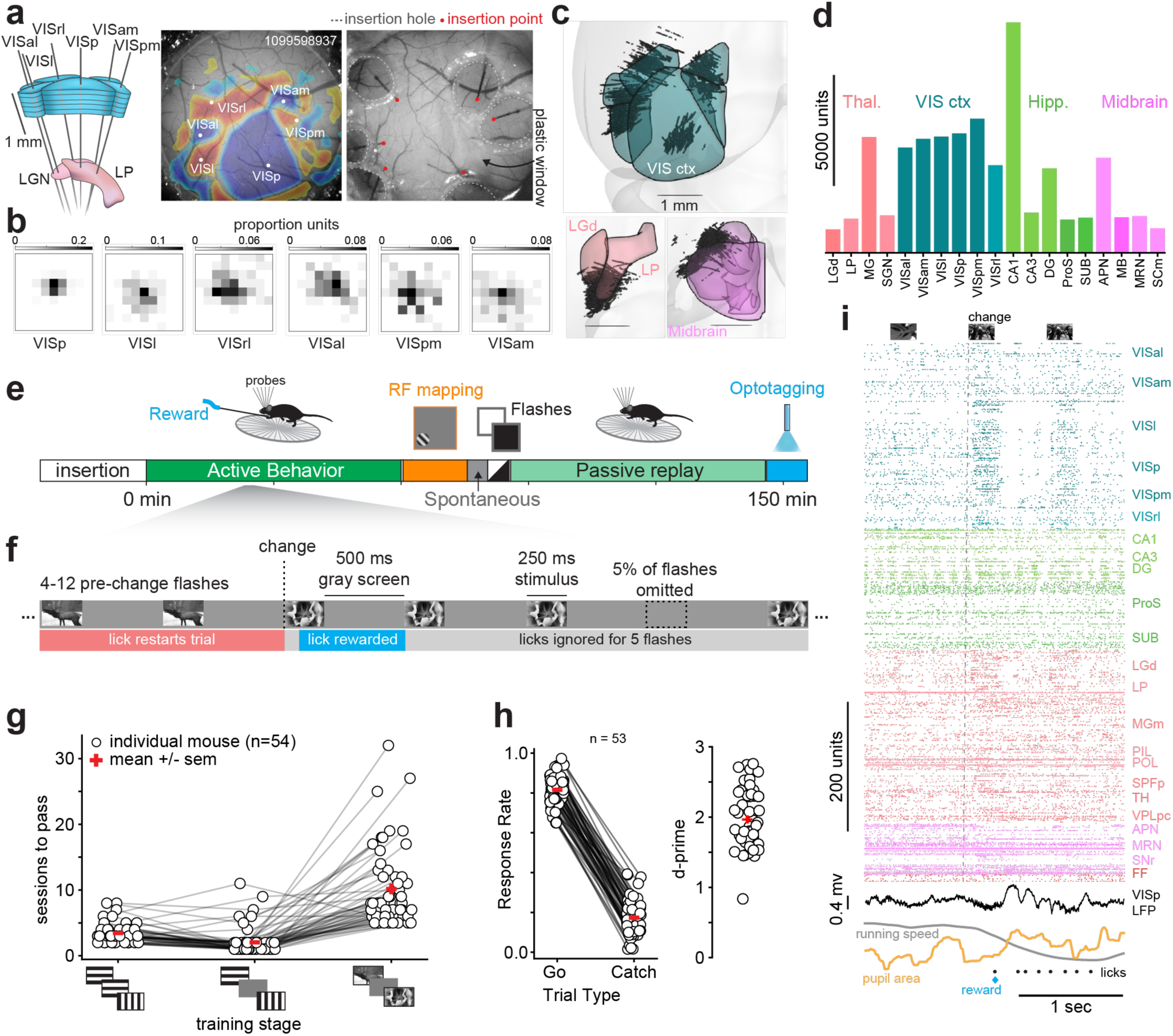
Survey of activity in visual cortex, thalamus and midbrain during visual change detection behavior. **a)** Left: Recording configuration targeting six visual cortical areas and thalamus with Neuropixels probes. Middle: Sign-map generated by ISI imaging identifying cortical area boundaries to guide probe insertions. White circles indicate probe insertion sites. Right: Image of brain surface during recording showing probe insertions (red dots) through the plastic insertion window. **b)** 2-D histograms showing distribution of population receptive field centers in each cortical area across experiments. **c)** Locations of recorded units superimposed on CCF volumes. Top: Unit locations relative to six visual cortical areas (blue). Bottom: Left, Units relative to thalamus (LGd, LP). Right, units relative to midbrain (SCm and MRN). For each panel, only units within the dorsal-ventral span of indicated regions are shown. Scale bars span 1 mm. **d)** Unit counts across all areas with at least 1000 units. **e)** Recording session timeline showing six experimental epochs. **f)** Timeline for example trial during change detection task. **g)** Number of sessions to pass the three stages of behavioral training for each mouse (white circles). **h)** Left: Response rates for change trials (go) and non-change trials (catch) across mice (white circles). Rates are calculated after combining trials across both recording days. Right: d-prime values calculated from go and catch response rates. Red crosses in (g) and (h) show mean ± sem across mice. Note one mouse from (g) failed to engage in the task during recording and was excluded from (h). **i)** Spike raster showing hit trial in example session. Dotted line denotes image change time. VISp LFP (black), running speed (gray), pupil area (gold), reward (blue) and licks (black dots) are shown beneath raster.

Each session comprised five experimental epochs: (1) active behavior, during which mice performed a visual change detection task for water reward; (2) passive stimulation with gabor stimuli to map receptive fields; (3) spontaneous activity with no visual stimulation; (4) full-field luminance changes; and (5) a “passive replay” epoch during which the mouse was presented with the same image sequence encountered during active behavior but with the lick spout retracted (Figure 1e). After these epochs, we used optotagging to identify cortical interneurons expressing ChR2 in *SST-cre;Ai32* or *VIP-cre;Ai32* mice (Figure 1e; Figure S2; 434 SST+, 27 VIP+ units).

During the visual change detection task, mice viewed a series of briefly presented natural images and received a water reward for licking when the identity of the image changed. Every trial, a given image was repeated 4-11 times before changing. Images were presented for 250 ms and separated by 500 ms of isoluminant gray screen. Approximately 5% of image presentations were randomly omitted, producing a gray interval of 1.25 s between image presentations (Figure 1f)^17,23^. During training, mice progressed through several stages (Figure 1g), reaching proficiency in the final task after 3-4 weeks of training (15.7 +/- 6.9 sessions, mean +/- std). Once proficient, mice licked with high probability on go trials (when a change occurred) compared to catch trials (when a change did not occur) (p=2.4e-10, Wilcoxon signed-rank test), resulting in high d-prime values (d-prime comparing go and catch responses = 2 +/- 0.4, mean +/- std, 53 mice; Figure 1h). In a previous study implementing this task with less stringent training criteria, behavioral analysis showed that mice used a combination of visual change detection and timing estimation to inform their lick decision^24^. Applying the same analysis, we found that the mice in this dataset overwhelmingly used visual cues to decide when to respond (see *Automated Training* in Methods).

In each experiment, we recorded spiking activity from ∼700 high-quality units (739 +/- 183, mean +/- std) and collected several streams of behavioral data including running speed, lick spout contacts, and videos of the eye, face and body (Figure 1i).

### Encoding of sensory and action variables

To characterize how units encoded visual stimuli, task events and motor output during the change detection task, we fit a general linear model (GLM) to the spiking activity of each unit during the active behavior epoch. This GLM seeks to explain the firing rate (25 ms bins) of a given unit by a linear combination of 14 time-dependent kernels: eight kernels for the eight unique natural images shown during a given session (visual features), as well as kernels for hits and misses (task features), image omissions, and finally, licks, running speed and pupil radius (action features; Figure 2a). The model fit captured 6.5% of the total variance on average across all areas and 11% for visual cortical units (all: 6.5 +/- 9.8%, mean +/- std; VIS: 11 +/- 11.8%; values reflect cross-validated performance; Figure S3a), consistent with similar efforts^5,12^.

**Figure 2:**
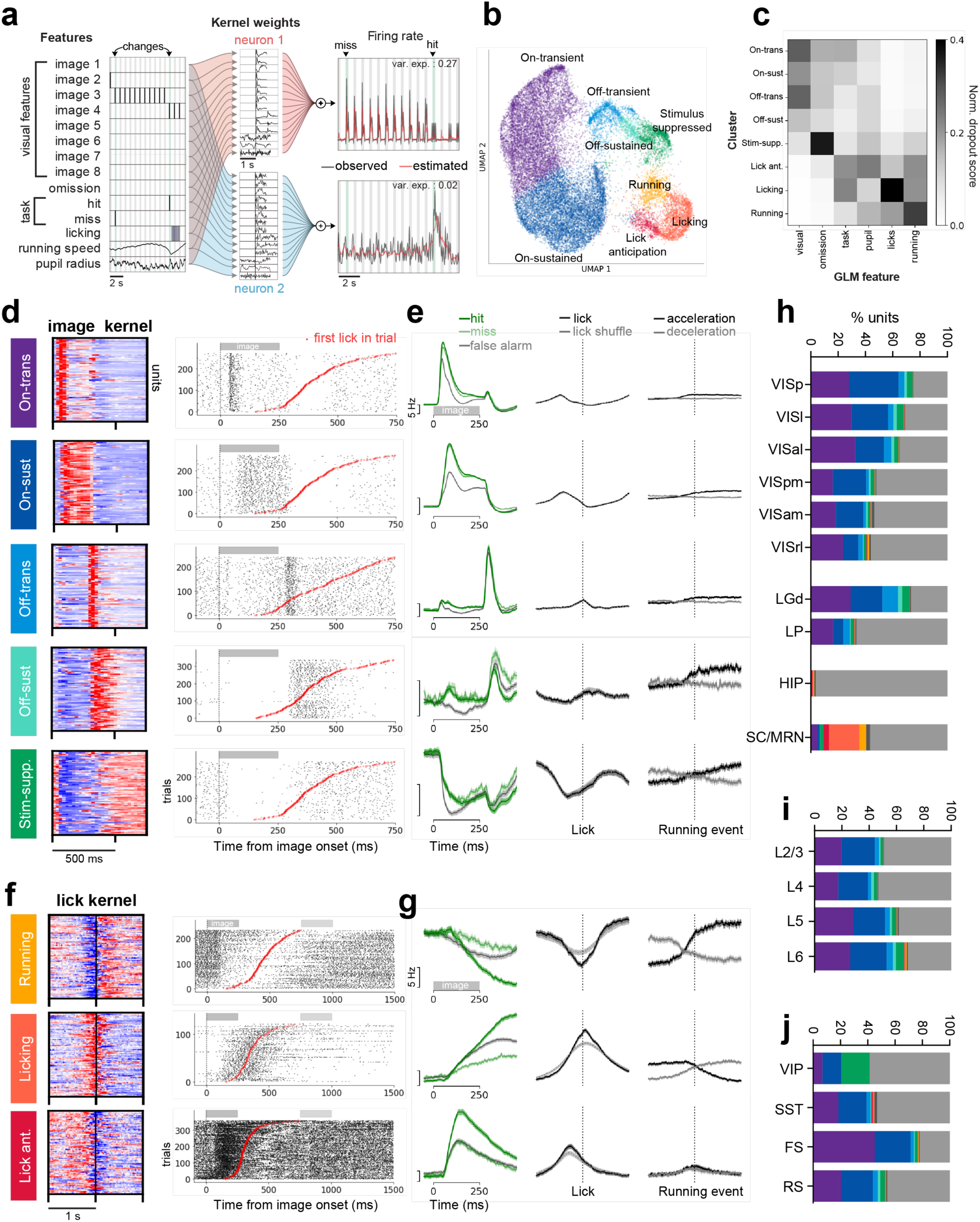
GLMs fit to single units reveal diverse sensory and action coding across visual cortex, thalamus and midbrain. **a)** GLM design showing how features for 14 kernels were constructed for a segment of an example session (left). Middle: kernel fits for two example neurons (red and blue shading). Kernels were convolved with features and summed to estimate firing rates. Right: observed (gray) and estimated (red) firing rates for example neurons. **b)** Uniform Manifold Approximation and Projection (UMAP) embedding of kernel fits for all recorded units colored by cluster label. **c)** Heatmap of normalized dropout score for kernels across clusters. Matrix is normalized so columns add to 1. **d)** Left: Preferred**-**image kernels for 100 example neurons in each sensory cluster. Right: spike raster for one example neuron in each cluster aligned to preferred image onset for non-change stimuli for which the mouse responded. Trials have been ordered by response time (red symbols). **e)** Population-averaged firing rates for each cluster aligned to image onset for hit, miss and false alarm trials (left), lick response (middle; response times were shuffled across trials and response was recalculated to produce gray trace) and running events (right; acceleration from stillness in black; deceleration to stillness in gray). Running events were taken from passive replay epoch to avoid correlation with licking. Shaded regions indicate SEM across units. **f,g)** same as (d,e) but for action clusters with kernels and rasters aligned to lick response times. Note that the time window is extended to show the motor response. **h-j)** For each region (h), cortical layer (i), and cortical cell class (j), percentage of units belonging to each cluster. Light gray bar indicates units for which the GLM did not explain a significant fraction of variance (<5%). Dark gray indicates units in three clusters excluded from further analysis.

To assess the degree to which units encoded sensory or action features, we compared the performance of the full model to one for which either the image kernels were removed (visual dropout) or the lick kernel was removed (lick dropout). The resulting dropout scores measure the unique contribution of the respective kernels to the variance explained for each unit. As expected, visual dropout scores were highest in visual cortical and thalamic regions, with VISp and lateral visual areas VISl and VISal scoring higher than anterior/medial visual areas VISrl, VISpm and VISam (Figure S3b). In the deep layers of the superior colliculus (SCm) and the midbrain reticular nucleus (MRN), ∼11% of units had high visual dropout scores (>1% change in total variance explained), while 29% had high lick dropout scores (Figure S3b), consistent with the premotor function of these midbrain structures^25,26^. We noted a rough dorsal/ventral topography of visual/motor encoding in SCm and MRN (Figure S3e,f). A small fraction (1%) of visual cortical units exhibited high lick-dropout scores; these units were more common in higher visual areas than VISp and were biased to deep cortical layers (Figure S3 c,d).

Two units may have similar dropout scores for a given feature but respond with different dynamics. To gain insight into the dynamics of visual, task, and motor encoding, we clustered units based on their temporal kernels (Methods). This yielded 11 clusters: 5 sensory clusters with distinct visual response dynamics, 3 action clusters, and 3 clusters which were either too small or too weakly associated with our GLM variables to be analyzed in detail (Figure 2b,c; kernels for all clusters are shown Figure S4). The eight clusters selected for further analysis were found to be robust by a bootstrapped classification method, which quantified the frequency with which a random forest classifier trained on a subset of the data correctly predicted the cluster labels for held-out units (Figure S5).

The 5 sensory clusters consisted of 1) an ‘on-transient’ cluster with a sharp response at stimulus onset, 2) an ‘on-sustained’ cluster with longer latency activity sustained throughout the stimulus presentation, 3) an ‘off-transient’ cluster with a sharp response at stimulus offset, 4) an ‘off-sustained’ cluster, and 5) a ‘stimulus-suppressed’ cluster, which was inhibited by the visual stimulus and showed gradual ramping activity during inter-stimulus intervals and stimulus omissions (Figure 2d,e; Figure S4, Figure S6a). Sensory encoding units were most common in visual cortex and thalamus but were present in every area recorded, with ‘on’ outnumbering ‘off’ units (Figure 2h). Midbrain sensory units were mostly on-transient (Figure 2h). Within cortex, stimulus-suppressed units were more common in the deeper layers and were particularly enriched in VIP+ interneurons identified by optotagging (Figure 2i,j; Figure S6b). As a population, sensory cluster units did not show significant lick-aligned activity relative to a shuffled control but were weakly modulated by running (Figure 2e). Midbrain neurons in the sensory clusters had more mixed encoding of sensory and motor information compared to visual cortical neurons (Figure S3g).

The three action clusters consisted of 1) a ‘running’ cluster with activity anticipating locomotion, 2) a ‘licking’ cluster with activity during licking bouts, and 3) a ‘lick-anticipation’ cluster whose activity preceded licking (Figure 2f,g; Figure S4). Action cluster units were most common in the midbrain (Figure 2h). In visual cortex, these units were rare and biased towards anterior/medial higher visual areas VISam and VISrl (Figure 2h) and deep cortical layers (Figure 2i).

In several aspects, lick-anticipation units in SCm/MRN appeared to reflect the decision to lick. These units were strongly active during stimuli that elicited licking (hits and false alarms; Figure S7a) and responded to stimuli at short latency (∼50 ms) during active behavior but not passive viewing (Figure S7a). Activity during the baseline period in these units was elevated before stimulus presentations that elicited false alarms (Figure S7a,c), and anticorrelated with response time (Figure S7g,i). Moreover, lick-anticipation units—but not lick units—showed ramping activity that increased with time from the last lick bout, mirroring an increased propensity for mice to commit false alarms at long inter-lick-bout intervals (Figure S7 a-f).

### Critical window for change detection decision

To investigate the chain of processing leading from image presentation to the lick response, we evaluated when sensory information (image identity and image change) and action information (the lick response) could be decoded in each area (Figure 3a). We used sensory cluster units to decode image-identity and image-change, and action cluster units to decode licking (see Figure S8a-c for comparison of decoding performance in sensory and action clusters). Because sensory and/or action units were sparse in some areas, we pooled units across sessions, creating pseudo-populations of 100 units for decoding (see Methods). Baseline activity was subtracted to emphasize stimulus-driven changes in activity. For each decoder, we selected training data to isolate the variable of interest (Figure 3b).

**Figure 3:**
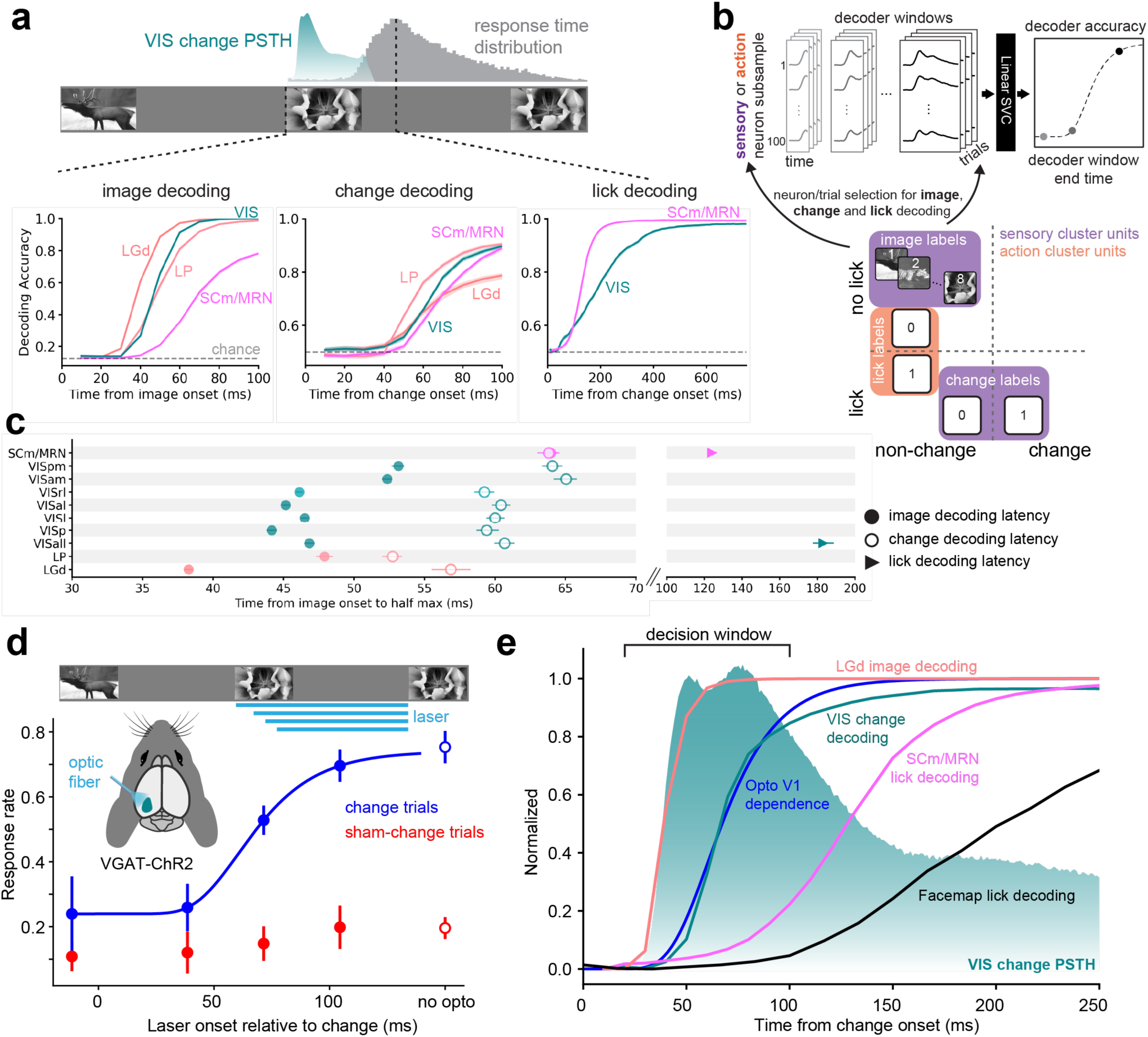
Decoding, behavior and perturbations reveal critical window for change decision. **a)** Top: schematic depicting relative timing of image change onset, population response in visual cortex (teal), and lick response time distribution across mice (gray). Bottom: Decoding time courses across areas for image (left), change (middle) and lick (right) decoding. Shaded regions give bootstrapped 95% CI of the mean across pseudo-populations. **b)** Schematic depicting decoding method. Each timestep on the decoding curve (top right) represents the performance of a linear decoder trained on activity in a window ranging from stimulus onset to that time point. Sensory cluster neurons were used for image and change decoding; action clusters were used for lick decoding. Pseudotrials were created by sampling 100 neurons for each area/cluster combination across sessions. Bottom: Diagram depicting which trial types were selected to decode image, change and licking. **c)** Mean decoding latency (time to half max) across neuron subsamples for image (filled circles), change (open circles) and lick (filled triangles) decoding. Error bars represent 95% CI of the mean across pseudo-populations. **d)** Top: schematic of VISp silencing experiment: an optic fiber delivered light to the cortical surface at four intervals relative to stimulus onset. Bottom: mean lick response rates (n=4 mice, 15 sessions) plotted as function of laser onset time for change trials (blue) and catch trials (red). Error bars indicate SEM. **e)** Selected decoding curves are normalized to maximum in 500 ms from change onset and plotted with the visual cortex change stimulus response to highlight the critical window during which mice compute change and decide to lick. Black curve represents decoding of the lick response from first 500 SVDs of a video of the face.

Image identity information was present shortly after the visual latency and its temporal propagation followed the visual hierarchy, with LGd leading (latency to half max: 38 ms) followed by VISp and the lateral areas (44-46 ms), then the medial areas VISpm and VISam (∼53 ms) and finally SCm/MRN (64 ms) (Figure 3a,c). Decoding image *change* was delayed relative to image identity (earliest change decoding: 53 ms; earliest imaging decoding: 38 ms), and was more compressed across areas (image change range, 12 ms; image identity range, 25 ms; Figure 3c). Notably, higher-order visual thalamic area LP led all areas in change decoding (latency 53 ms), followed by LGd/VISp and lateral areas VISl/VISal (∼60 ms), and lastly the medial areas VISpm, VISam together with SCm/MRN (∼64 ms). LP continued to lead VISp after matching units from the two regions across sessions (Figure S8d,e; LP latency: 51.4 ± 3.2 ms, VISp latency: 58.4 ± 4.1 ms; mean ± standard deviation; p = 1.1e-15, Wilcoxon signed-rank test). SCm/MRN was the only area lacking a delay between change and image decoding latencies.

Licks could be decoded reliably from SCm/MRN action neurons just ∼120 ms after image onset, placing an upper bound on the time window during which image information is used by mice to inform the decision to lick (Figure 3a,c). Decoding licks from action neurons in visual cortex lagged the midbrain by about 50 ms, possibly reflecting an efferent signal. Action neurons were too sparse in individual visual cortical or thalamic regions to decode licking by this method.

To causally test when information in the visual cortex was used by the mice to detect image change, we optogenetically silenced VISp in VGAT-ChR2 mice^27,28^, starting at various timepoints relative to the image change and continuing to the end of the response window (n=4 mice, 15 sessions; Figure 3d). When VISp silencing started prior to the change image presentation, the behavioral response rate was potently reduced. However, behavior was not impacted when VISp silencing began 100 ms after the image change, indicating that mice required no more than 100 ms of VISp activity to reliably detect image changes. At an intermediate silencing timepoint, change detection was partially disrupted. This time course of VISp dependence closely resembled the time course for change decoding in visual cortex (Figure 3e).

The mouse’s lick response is an upper bound on the time of action initiation, since orofacial movements including jaw opening precede licking. To further constrain the time when overt motor output might influence neural activity through reafference, we trained a decoder to identify lick bouts from video data of the face (see Methods section *Facemap lick decoding*). This provided a much faster estimate of the behavioral response time than lick latency (face decoder latency: 201 ms, median first lick time: 424 ms), but still lagged midbrain lick decoding by ∼75 ms (time to half max for face decoder: 201 ms, midbrain decoder: 123 ms) (Figure 3e).

Given these bounds, we operationally defined a decision window extending from 20-100 ms after image presentation, starting just before the first stimulus driven spikes in visual thalamus, ending just before an inflection point in midbrain lick decoding and well before the time at which licking could be reliably decoded from facial movements (Figure 3e).

### Modulation by change and task-engagement across areas, layers and cell types

Having identified the window during which visual system activity could inform the decision to lick (20-100 ms after stimulus onset), we next investigated which cell populations were most modulated by behaviorally relevant stimuli (image changes) and task-engagement during that interval. For this analysis, we were most interested in modulation of the sensory response and focused our analysis on stimulus-evoked activity in sensory cluster units. To quantify the degree to which individual units were modulated by image changes or engagement, we defined two indices: a ‘change modulation index’ comparing responses to image changes with responses to the stimuli immediately preceding changes (‘pre-change’ stimuli; Figure 4a), and a ‘state modulation index’ comparing responses to change stimuli shown during the behavioral task with those during passive replay (Figure 4a).

**Figure 4:**
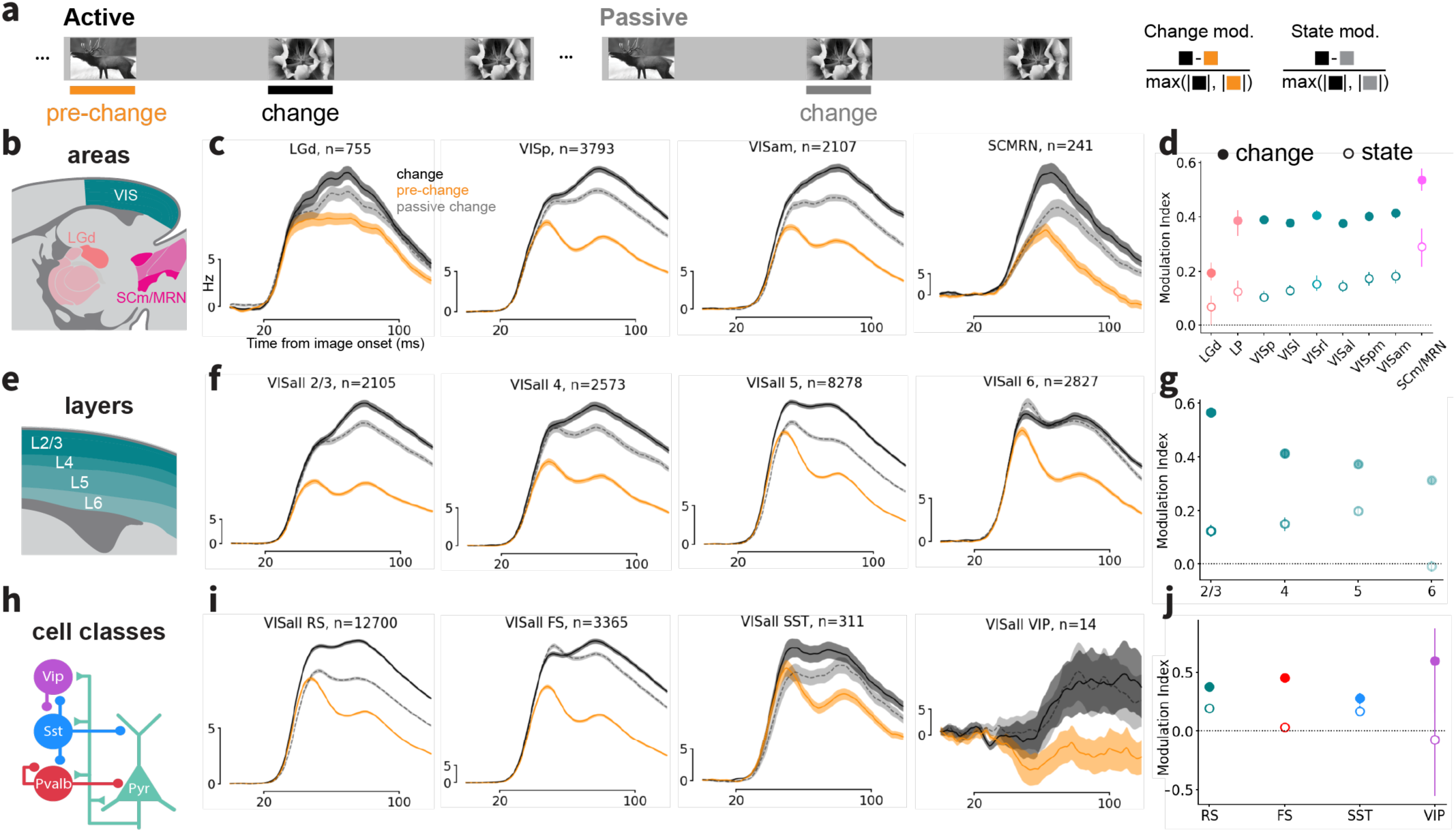
Modulation by image change and task engagement across the visual system. **a)** Schematic depicting quantification of change and state modulation. The change modulation index compares responses to change and pre-change stimuli during the active behavior epoch. State modulation compares responses to change stimuli during the active and passive epochs. Modulation indices were computed on evoked activity for units in sensory clusters during the decision window (20-100 ms after image onset). **b)** Diagram depicting areas for which change/state modulation is analyzed in (c) and (d). **c)** Population PSTHs for active change (black), active pre-change (orange) and passive change (gray). Y-axis scale bar indicates firing rate of 5 Hz. X-axis scale bar indicates the decision window used to compute metrics in (d). Shaded region indicates SEM. **d)** Change (solid symbols) and state (open symbols) modulation indices for nine areas spanning visual system. Symbols indicate median across population and error bars indicate bootstrapped 95% CI of the median. **(e-g), (h-j)** Same as b-d but for cortical layers (e-g) and cell classes (h-j). Layer and cell-class data were pooled across visual cortical areas.

Across areas, change modulation was stronger in visual cortex than LGd, but even stronger in SCm/MRN units (Figure 4b-d, Figure S9a,c; comparing black and gold curves). Within cortex, layer 2/3 units were more modulated than the lower layers (Figure 4e-g, Figure S9c), while cortical SST+ units were less modulated than the other cell classes (Figure 4h-j, Figure S9c).

State modulation increased across the thalamocortical visual hierarchy, and SCm/MRN units were again most strongly modulated (Figure 4b-d, Figure S9a,d; comparing black and gray curves). Across cortical layers, layer 6 was the only layer not significantly modulated by task engagement, while layer 5 showed the strongest modulation (Figure 4e-g, Figure S9d). Across cortical cell types, regular spiking (RS) and SST units were more strongly modulated by task engagement compared to fast spiking (FS) units. VIP units were highly variable (Figure 4h-j, Figure S9d). These findings could not be explained by differences in running speed across active and passive epochs (Figure S10).

In summary, modulation by both image change and task-engagement depended on position along the visual hierarchy. RS cells exhibited a strong preference for change stimuli and a weaker preference for the active behavioral context, but this effect varied across layers, with layer 6 units showing little modulation (Figure S9b). Units in the midbrain nuclei SCm/MRN showed the strongest change- and task-engagement modulation, indicating that sensory responses in these areas are potently gated by the behavioral relevance of the stimulus.

### Stimulus novelty modulation arises in visual cortex

The change modulation index above reveals how units respond to ‘contextual novelty’ over short time scales (∼10 seconds). We designed our study also to investigate how long-term familiarization with an image set shapes neural activity relative to stimuli the mouse has never seen (‘absolute stimulus novelty’). Throughout training, mice encountered only one set of eight images (either image set ‘G’ or image set ‘H’; Figure 5a,b), after which we performed two recordings from a given mouse on consecutive days. On one recording day, the mouse performed the change detection task with the familiar image set used during training. For the other recording day, we used a novel image set, consisting of six new images the mouse had never seen and two ‘holdover’ images from the familiar set (purple background in Figure 5b). For each cell population, we computed a novelty modulation index based on population responses to these two image sets, excluding the holdover images (Methods).

**Figure 5:**
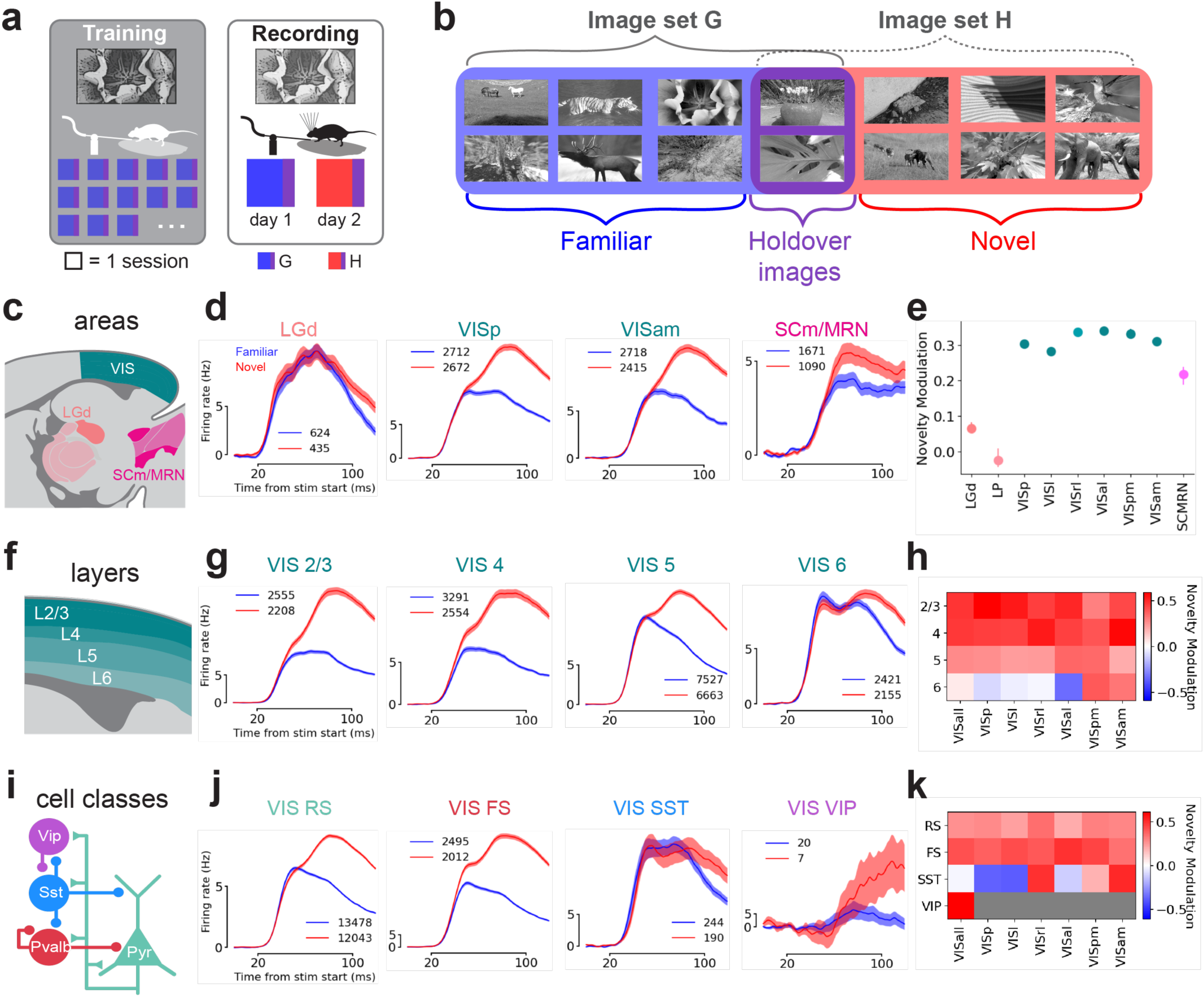
Modulation by stimulus novelty across the visual system. **a)** Diagram depicting the history of image-set exposure for a typical mouse during training and recording days. Each colored square indicates one behavioral session. During training (left) the mouse encounters only one image set (G for this example mouse). During recording (right), the mouse is presented with the training image set for one recording day and a novel image set (H in this example) for the other recording day. **b)** G and H image sets each comprised eight images. Two images were shared across the two image sets (‘holdover images’, purple background). To compare familiar and novel image responses, only the six images unique to each set were used. **c)** Area diagram. **d)** Population PSTHs for familiar (blue) and novel (red) images across four areas. Shaded region indicates SEM. **e)** Novelty modulation indices for nine areas spanning visual system. Novelty modulation was computed on evoked activity during the decision window (20-100 ms from image onset). Error bars indicate bootstrapped 95% CI for median. **(f-h), (i-k)** Same as c-e but for cortical layers (f-h) and cell classes (i-k). Note that in (k) novelty modulation indices for VIP+ units could not be calculated for individual areas due to their limited number.

Responses in visual thalamic regions LGd and LP were minimally influenced by image novelty, whereas all visual cortical areas showed strong novelty-enhancement (Figure 5c-e). Midbrain structures SCm/MRN had an intermediate preference for novel images (Figure 5c-e). These results were qualitatively similar across the active and passive conditions as well as change and non-change stimuli (Figure S11a,b).

Within cortex, novelty modulation was strongest in the upper layers, with less modulation in layer 5 and little to no novelty enhancement in layer 6, particularly in the lateral/posterior visual areas (Figure 5f-h, Figure S11e). This lack of modulation in layer 6, the main source of cortical feedback to primary visual thalamus, is consistent with the weak novelty modulation observed in LGd.

Across cortical cell types, RS, FS and VIP population activity was strongly increased by image novelty, whereas the SST population had similar activity for novel and familiar images (Figure 5i-j). Prior calcium imaging experiments showed that SST cells in VISp and VISl prefer familiar images^12^. To reconcile these findings, we split SST units by cortical area. Consistent with previous studies, we found that SST units in VISp as well as lateral areas VISl and VISal were indeed familiar-image preferring (Figure 5k, Figure S11d). In contrast, SST units in anterior/medial areas VISrl, VISpm and VISam preferred novel images (Figure 5k; Figure S11d).

Given the lack of novelty modulation in thalamus, we next considered whether the novelty enhancement we observe in cortex emerges through cortical interactions. If so, we should see evidence for a recurrent cortical origin in the temporal dynamics of novel image responses: specifically, novelty modulation should arise after the first sweep of activity up the visual hierarchy.

Indeed, we found that the first wave of image-evoked activity in LGd and cortex was not different between familiar and novel images (Figure 6a,b) and flowed along the anatomical hierarchy as expected (Figure 6c). In contrast, novelty modulation manifested as a clear second peak in the cortical PSTH ∼60 ms after stimulus onset (Figure 6b,d) and only appeared in visual thalamus at the end of the decision window (∼100 ms, Figure 6a,d). In RS cells, the disparity in modulation between the early and late response was most dramatic, with the initial peak slightly preferring the familiar image set (Figure 6b). Importantly, the time course of novelty modulation in cortex was the same for mice trained on either image set G or H (Figure S11f). Comparing familiar holdover images to novel images on the novel recording day revealed similar delayed modulation for RS and FS units (Figure S11d).

**Figure 6:**
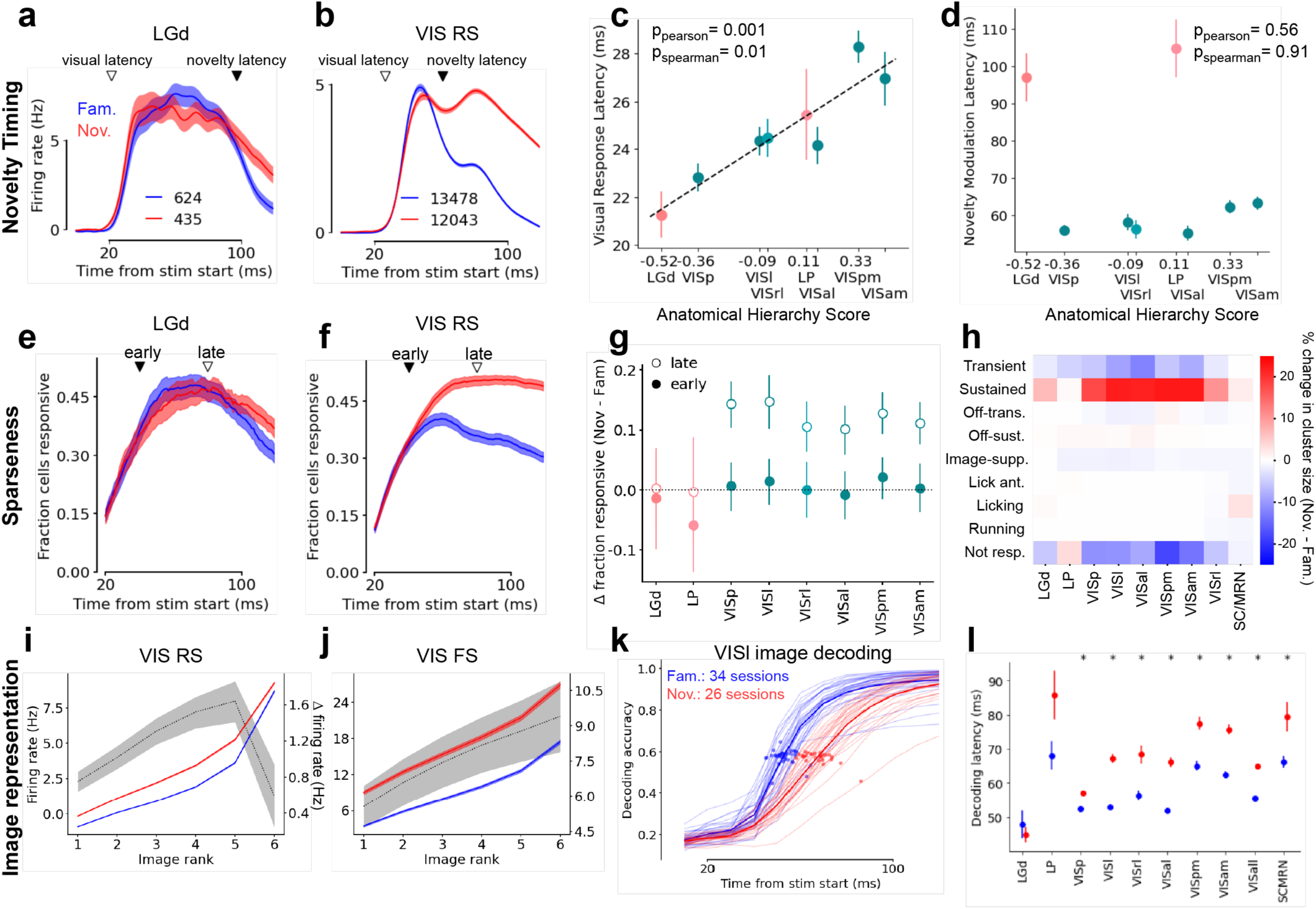
Novel images recruit slow, dense population responses in cortex. **a)** LGd population PSTH for familiar (blue) and novel (red) non-change stimuli. Shaded region indicates SEM. Open arrowhead indicates estimated visual latency. Closed arrowhead indicates estimated novelty preference latency. **b)** Same as (a) for visual cortical RS units. **c)** Visual response latency plotted against visual hierarchy position for cortical and thalamic areas. Note that for comparison to the novelty modulation latency in (d), visual latency is calculated on the population response. **d)** Novelty modulation latency plotted for same areas as (c). Error bars for (c,d) indicate standard deviation of the bootstrap distribution of means (1000 iterations). **e)** Fraction of LGd units responsive over time for familiar and novel images. Closed arrowhead indicates the midpoint of the early response window (20-60 ms after stimulus onset). Open arrowhead indicates the late response window (60-100 ms). **f)** Same as (e) but for VISp units. **g)** Difference in fraction of units responsive to novel (closed symbols) and familiar (open symbols) images for early and late response windows in (e,f). Error bars indicate 95% CI of the mean. **h)** Change in cluster membership for units recorded in novel sessions versus familiar sessions. **i)** Mean tuning curves of visual cortical RS units for familiar and novel stimuli. For each unit, image responses were ranked before averaging. Dotted gray line indicates difference between novel and familiar curves. For red and blue curves, shading indicates SEM. For gray curve, shading indicates 95% bootstrapped CI. **j)** Same as (h) for visual cortical FS units. **k)** Image decoding time courses for VISl units during 34 familiar and 26 novel sessions. Light traces indicate individual sessions. Dark traces are means across sessions. Symbols indicate time to half-max for each session. **l)** Mean image decoding latency across areas. Error bars indicate SEM across sessions. * indicates p<0.05 for Wilcoxon rank-sum test after correction for multiple comparisons.

The delayed novelty amplification we observed in cortex could reflect an increase in the firing rate of neurons that were already responsive in the early time window and/or a recruitment of a new population of neurons with longer latency responses. We quantified the fraction of cells with significantly elevated novel-image responses in the first half (20-60 ms from stimulus onset; ‘early’ epoch) and second half (60-100 ms; ‘late’ epoch) of the critical decision window. In the early epoch, a similar fraction of RS cells responded to novel and familiar images, but in the late epoch 10-15% more cells responded to the novel image set (Figure 6f). This pattern was observed in all cortical areas (Figure 6g). In visual thalamus, neither epoch showed a recruitment of more units by novelty (Figure 6e,g).

Examining sensory cluster membership for familiar and novel recording sessions provided additional insights into the expanded cortical population responding to novel stimuli. In novel sessions, the fraction of ‘on-sustained’ units increased, whereas ‘on-transient’ and ‘non-responsive’ units decreased (Figure 6h). Changes in the other sensory or action clusters were negligible (Figure 6h).

Taken together, the reduced sparseness in the late response and the change in cluster membership for novel stimuli suggest that many neurons which were not previously responsive to familiar images are recruited at long latency by novel images. What consequences do these changes have on how images are represented in visual cortex? We found that the impact of novelty on image tuning was cell-type specific, consistent with previous results in primates^29^. Tuning of RS units in visual cortex broadened due to amplification of less-preferred stimuli (Figure 6i), whereas modulation in FS units monotonically scaled with image preference (Figure 6j). Despite these changes, familiar and novel images were decoded with similar accuracy across all visual areas (Figure S12a), although decoding of novel images lagged familiar images, particularly in the higher visual areas (Figure 6k,l). Thus, familiarization sharpened both the temporal dynamics and stimulus tuning in RS cells, producing similar decoding accuracy with fewer spikes and faster latencies.

In four mice, we investigated the time course of novel image familiarization across days (Figure S12b). Consistent with results from a previous calcium imaging study^12^, we found that the magnitude of the response to novel images returned to familiar levels after one session of exposure (Figure S12c,d). However, the time course of the response after one session was still slower than that for familiar images (Figure S12c,e,f).

### Perturbing behavior with novel images reveals change detection strategy

Given the dramatic changes in neural representation of novel versus familiar images, we next investigated whether behavior with the novel image set could yield clues about the computational strategy mice used to solve the task. We considered two strategies, which, given the impact of novelty on image responses, make very different behavioral predictions for the two familiar holdover images (purple images in Figure 5b) during novel and familiar sessions.

In the first strategy, mice remember the identity of the last image shown until the next image presentation. When the next image comes, they read-out the identity of that image and compare it to the remembered image, licking for a mismatch. To model this ‘image comparison’ strategy, we trained a one-vs-all image decoder on activity during the 20-100 ms decision window to determine whether each image presentation was the same as or different from the previous image. The judgement of the decoder was quantified by the decoder confidence, given by the distance of each image response from the decoder boundary learned from training data (magenta curly bracket in Figure 7a).

**Figure 7:**
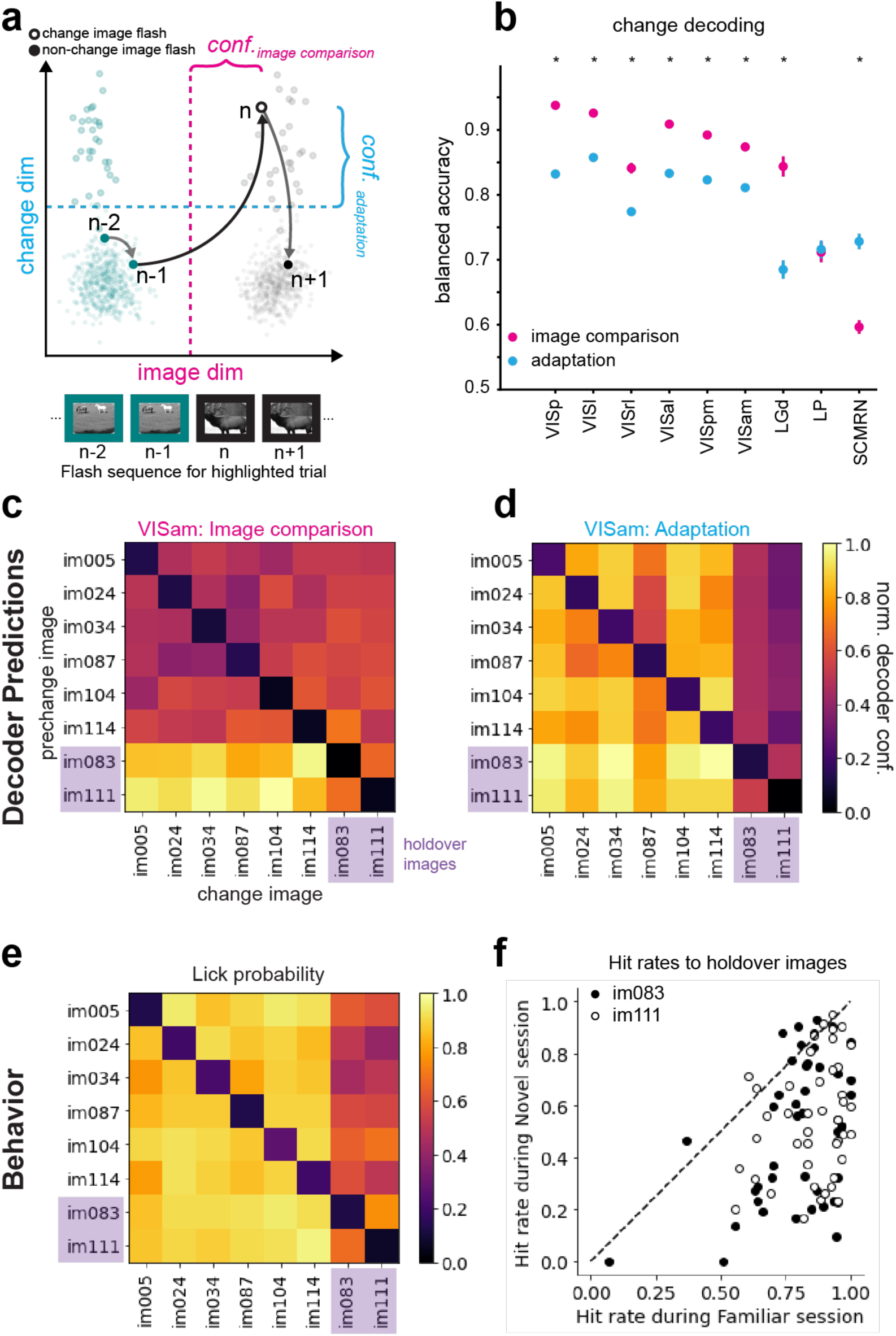
Behavior during novel day reveals change computation strategy. **a)** Plot illustrating two possible strategies for detecting image change. Individual dots are population responses in visual cortex to stimulus presentations plotted in two dimensions maximizing separation between images (x axis) and between change and repeat stimuli (y axis). Responses to two example stimuli are shown in teal and black. Solid symbols are repeat presentations and open symbols are change stimuli. Bold symbols show responses from one example trial when the image transitioned from the teal to black image. Magenta line indicates boundary for decoding image identity (‘image-comparison strategy’). Cyan line indicates the boundary for decoding change (‘adaptation strategy’). Distance from these boundaries measures the confidence of the respective decoder. **b)** Change decoding accuracy for logistic regression using confidence from either the image-comparison decoder (magenta) or the adaptation decoder (cyan). Error bars indicate SEM. * indicates p<0.05 for Wilcoxon signed-rank test after correction for multiple comparisons. **c)** Heatmap matrix showing the normalized mean decoder confidence across sessions for the image comparison decoder for each image transition during the novel day. Holdover images are the last two rows/columns. **d)** Same as (c) but for the adaptation decoder. **e)** Mean lick probability across sessions for each image transition during the novel day. **f)** Response probability for the two holdover images during the familiar and novel sessions. Each symbol represents the response rate to one of the images (open symbols for image 111; closed for image 83) for one session.

In the second strategy, the mice do not explicitly remember the identity of each image presentation. Instead, they take advantage of change modulation (Figure 4) to discriminate between change stimuli and non-change stimuli (i.e. repeats). For convenience, we refer to this as the ‘adaptation strategy’, though change modulation may not entirely arise from low-level adaptation mechanisms. To model the adaptation strategy, we trained a decoder to classify individual image presentations as either change or non-change and quantified the decoder confidence for each image presentation (cyan curly bracket in Figure 7a).

Both strategies successfully identified image changes well above chance across all areas tested (Figure 7b). The image comparison strategy was more accurate in LGd and visual cortex, whereas the adaptation strategy was more accurate in SCm/MRN (Figure 7b). However, breaking out decoder confidence for each image transition revealed starkly different predictions by the two strategies. The image comparison decoder predicts a benefit for detecting changes from familiar to novel images (first six columns of bottom two rows in Figure 7c), leveraging the separation of novel and familiar responses in population space. In contrast, the adaptation decoder predicts a deficit in detecting transitions to familiar holdovers (rightmost columns in Figure 7d), confusing response attenuation due to adaptation with that due to familiarization. Strikingly, we found that mice exhibit this same deficit (Figure 7e), responding to transitions to familiar holdover images at a much lower rate than transitions to novel images. This was not due to a general deficit in detecting changes for these stimuli, as mice responded at high rates for these same images when they appeared in the familiar session (Figure 7f). The same deficit was present in mice regardless of whether they trained on the G or H image sets, indicating that it was specific to the novel context rather than to a particular image set (Figure S13a, b).

Examining false alarms provided further evidence that the adaptation strategy better explained mouse behavior. The adaptation model predicted that mice should commit false alarms at a higher rate for novel images (diagonal of 7d), a pattern of response we also found in mouse licking behavior (Figure S13c). In addition, in all areas except LGd, the adaptation decoder better predicted trial-by-trial licking compared to image comparison (Figure S13d), particularly in higher visual areas and SCm/MRN. Finally, we found little evidence for a persistent representation of image identity during the inter-stimulus gray period (Figure S13e) as might be predicted by the image-comparison strategy, though this representation could be stored in brain regions not sampled in this study.

## Discussion

This study presents a large-scale database of spiking activity from mice performing a visual change detection task. Our findings define the time window during which visual cortical activity influences decision-making and illuminate the computational strategy used by the mice. Through dense sampling, we identify area-, layer-, and cell type-specific spiking dynamics during visual behavior and their striking modulation by novel stimuli.

### Dense neuronal sampling reveals response diversity across areas, layers and cell types

By recording across the visual system, from early sensory regions (LGd and VISp) to midbrain premotor areas (SCm and MRN), we were able to map how sensory and motor information is distributed across the visual hierarchy during visually-guided behavior. We found action-encoding units even in early visual cortical areas, supporting an emerging view that information about upcoming movements is widely distributed across the brain^5,30–32^ Action encoding in visual cortex was more prevalent in higher visual areas than VISp and was concentrated in deep cortical layers. Conversely, we found sensory neurons in SCm and MRN, midbrain regions associated with motor output^25,33^. Sensory neurons in these structures responded to images at short latency, and their activity could be used to decode the identity of images with high accuracy. Such distributed sensorimotor representations may play a critical role in active sensation, during which stimulus and movement information must be integrated to predict the consequences of a particular action^34^.

Recording from thousands of units in each visual cortical area allowed us to resolve differences across cortical layers and cell-types. We highlight two of these findings: 1) cortical layer 6 maintains stable sensory responses, and 2) SST+ inhibitory neurons show area-specific novelty modulation.

We identified cortical layer 6 as being less modulated by both task-engagement and stimulus novelty compared to the other layers. Since layer 6 provides the primary feedback projection to LGd, this lack of modulation is consistent with the similarly low levels of modulation by these factors in LGd. This compartmentalization may allow layer 6 and LGd to maintain a more veridical representation of the sensory environment that remains stable across task-engaged states and stimulus experience history. A previous study described heterogeneous effects of behavioral state on layer 6 neurons, but this modulation was related to major changes in behavioral state, ranging from sleep to high arousal, and was not compared to modulation in other layers^35^. Layer 6 is known to influence the gain of activity in the LGd and other cortical layers in a manner that strongly depends on the stimulus-context of visual input^36–40^. The lack of strong task-engagement and novelty modulation could allow layer 6 to implement this stimulus-specific control independent of cognitive and behavioral factors.

Using optotagging we identified SST+ inhibitory interneurons across all 6 visual cortical areas, which allowed us to examine whether SST+ cells had area-specific functional properties. Consistent with previous results using calcium imaging of SST+ cells in VISp^12^, we found that optotagged SST+ neurons in VISp and the lateral higher visual areas (VISl, VISal, VISrl) preferred familiar to novel images. On the other hand, SST+ neurons in the anterior/medial cortical visual areas (VISrl, VISam, VISpm) preferred novel images, indicating region-specific functional heterogeneity. This differential novelty modulation may relate to differences in recurrent connectivity across areas, as previous studies have identified stronger recurrent input to SST+ cells in medial versus lateral visual cortical areas^41^. Since our findings suggest that novelty modulation arises through recurrent cortical interactions, the enhanced recurrent connections onto SST+ cells in medial visual areas could underlie the novelty modulation observed in these cells.

### Time window for change detection behavior

A key aspect of our analysis was to define the critical time window (20-100 ms post-stimulus) during which stimulus change information is processed by the visual cortex to inform behavioral choices. We found a tight temporal correspondence between image change decoding from cortical activity and the behavioral effects of optogenetic silencing of primary visual cortex. This short window is consistent with previous studies that evaluated when visual cortex is necessary for stimulus detection and discrimination^42,43^. Intriguingly, stimulus changes could be decoded earliest in the higher-order visual thalamic area LP (mouse pulvinar)—even earlier than VISp—suggesting that LP may be a critical node in the change detection computation. Though a lag in the decoding time course between two regions does not necessarily indicate a processing lag (due, for example, to the impossibility of knowing the relevant population size for a given region), this result is consistent with LP’s proposed role in amplifying prediction errors in cortical layer 2/3 when unexpected stimuli are encountered^44^. Similarly, we find layer 2/3 is particularly sensitive to image changes, providing additional support for its role in representing prediction errors^45^. The rapid propagation of image change signals through this thalamocortical network, culminating in midbrain motor preparation within 120 ms, reveals the remarkable efficiency of this sensorimotor transformation.

Within this critical time window, we identified the computational strategy used by mice to perform the task, distinguishing between an “adaptation” versus “image comparison” computation. Using decoders trained on neural activity, we found that these two strategies could similarly detect image changes on average. However, the decoders made divergent predictions regarding the pattern of change detection performance in sessions with novel versus familiar images (Figure 7). Mouse behavior clearly supported the “adaptation” strategy over the “image comparison” strategy, as evidenced by specific deficits in detecting changes to familiar holdover images when presented in a novel context. Moreover, we found little evidence for an image-specific memory trace in visual cortical activity during the inter-stimulus gray periods (Figure S13e).

The adaptation strategy may be computationally advantageous because it requires only one decision boundary to classify ‘change’ versus ‘non-change’ stimuli, whereas distinguishing each image from the others would require up to eight boundaries. Additionally, this strategy doesn’t require maintenance of image templates in memory through energetically costly persistent spiking. Instead, adaptation to repeated stimulus presentations is an efficient solution that likely relies on short-term plasticity mechanisms including synaptic depression^46,47^.

### Novelty expands the active cortical population through recurrence

During the short 80 ms decision window following stimulus presentation, we observed distinct waves of activity in the visual cortex. The first wave (20-60 ms after stimulus onset) represents a feedforward sweep of activity up the visual hierarchy and is minimally affected by stimulus novelty. In contrast, a second wave of activity occurring between 60-100 ms is strongly modulated by stimulus novelty. This second wave likely reflects recurrent cortical processing, given the lack of novelty modulation in LGd yet prominent modulation in all cortical areas. During this window, novel images engage an expanded set of cortical neurons, recruited largely from a reservoir of ‘non-coding’ cells (Figure 6). VIP+ interneurons are also activated by novel images during this late window (Figure 5j), suggesting that VIP+ cells, perhaps through their inhibition of SST+ cells and consequent disinhibition of excitatory cells, may gate the recruitment of this expanded novelty-responsive population^12,48–51^.

The expansion of the number of responsive neurons with novelty may be computationally beneficial, providing cortex with the capacity to adapt to new stimulus statistics based on experience without disrupting existing representations^52,53^. The delayed novelty preference in cortex is not simply relayed to midbrain premotor regions (SCm/MRN), which show less novelty modulation. This may reflect the source of cortical input to these regions (primarily layer 5, which shows weaker novelty modulation than the upper layers) and/or integration with other non-novelty-modulated inputs. Together these results suggest a specialization of cortex for adaptive sensory representation via cortical recurrence, while premotor midbrain regions are tuned for behavioral relevance, evident from their strong stimulus change and state modulation.

### A database for mining and modeling

Overall, this dataset represents the most comprehensive characterization to date of spiking activity in the mouse visual system during a visually guided behavior and provides a rich resource for further data mining and modeling. To facilitate these efforts, we have packaged all spiking, LFP and behavior data (including the entire training history for each mouse) into Neurodata Without Borders (NWB) files and deposited them on public cloud storage platforms (Figure S1, Table 1). In addition, we provide a collection of tutorials and technical resources to help potential users understand, access and analyze this dataset (Table 2). Indeed, several studies have already utilized the database we describe to explore a variety of topics including omission responses across brain regions^54^, the impact of novel images on the representation of familiar images^55^, and how dynamics across the visual hierarchy approach criticality^56^.

Looking forward, this database of spiking activity opens numerous avenues for future exploration. Given the simultaneous recordings from six visual cortical areas, this dataset is an ideal substrate for examining multi-regional interactions^14^ and spike propagation through hierarchical networks^4^. Our dataset’s standardized and comprehensive nature—spanning multiple areas, cortical layers, and cell types during behavior— makes it an unprecedented resource for data-intensive models, including detailed biophysical models^57,58^, cell type-specific population models^59,60^, and artificial neural network foundation models^57,61,62^. The full impact of this database will emerge through extensive community engagement in mining, analysis, and modeling efforts.

## Acknowledgements

We thank the Allen Institute founder, Paul G. Allen, for his vision, encouragement, and support. We thank the Transgenic Colony Management, Animal Care, and Laboratory Animal Services teams at the Allen Institute for caring for the mice in this study. Funding for this project was provided by the Allen Institute. Anton Arkhipov and Omid Zobeiri were further supported by the National Institute of Mental Health of the National Institutes of Health under award no. U01MH130907. The content is solely the responsibility of the authors and does not necessarily represent the official views of the National Institutes of Health.

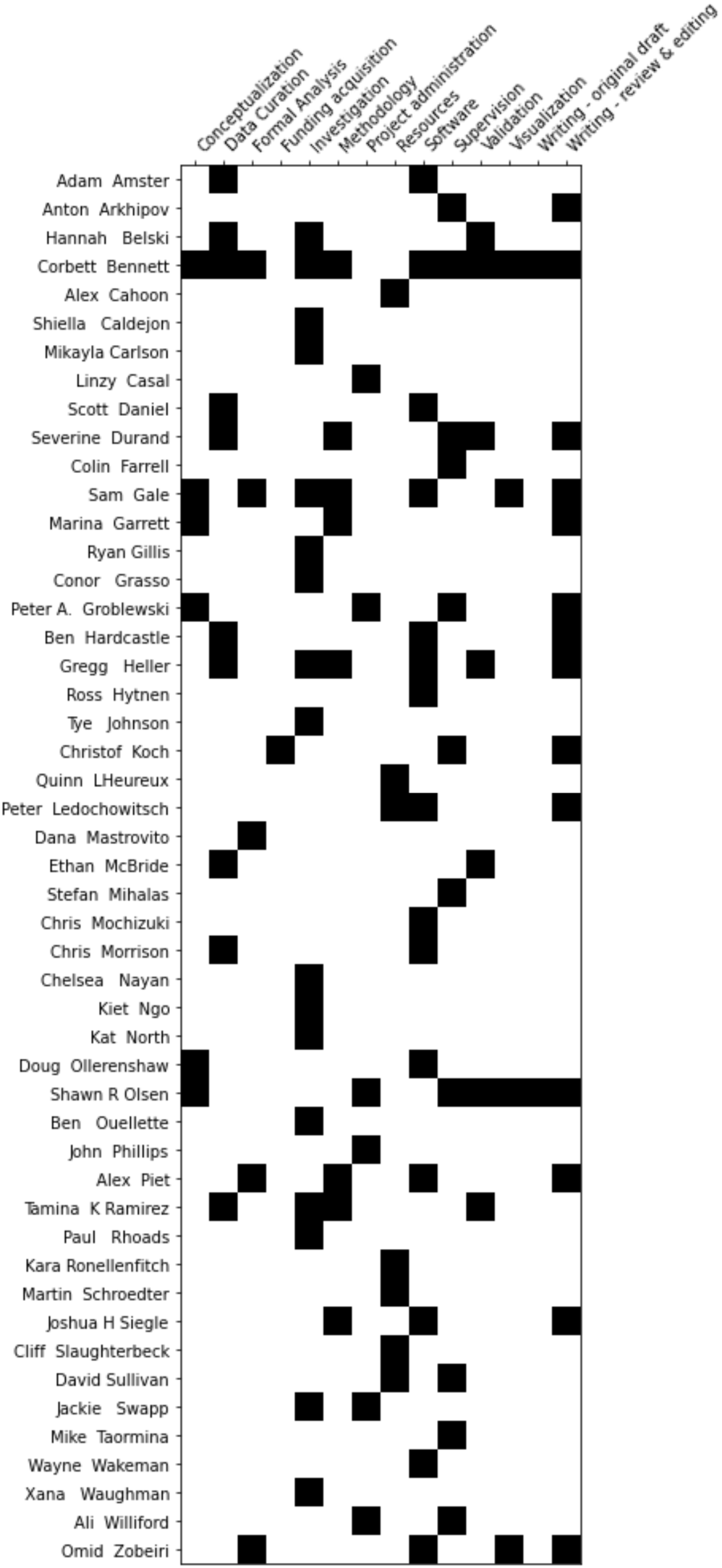

## Methods

### Mice

Mice were maintained in the Allen Institute animal facility and used in accordance with protocols approved by the Allen Institute’s Institutional Animal Care and Use Committee. The main dataset consisted of electrophysiology experiments from 27 C57BL/6J mice (15 males, 12 females), 31 Sst-IRES-Cre; Ai32 mice (16 males, 15 females), and 23 Vip-IRES-Cre; Ai32 mice (16 males, 7 females). Optogenetic perturbation experiments were performed in four VGAT-ChR2 mice (1 male, 3 females). Four-day electrophysiology experiments were performed in five Vip-IRES-Cre; Ai32 mice (3 male, 2 females). Following surgery, all mice were single-housed and maintained on a reverse 12-hour light cycle in a shared facility with room temperatures between 68° and 72°F and humidity between 30 and 70%. All experiments were performed during the dark cycle. During behavior training, mice were water restricted to 85% of their initial body weight with ad libitum access to food.

### Surgery

Mice received a headpost and cranial window as previously described^1,2^. Briefly, mice (aged 54 ± 6 days; mean ± standard deviation) were deeply anesthetized with isoflurane prior to exposing the skull. An incision was made to expose the skull and the head was levelled with respect to pitch, roll and yaw. The headframe was positioned to center the well at 3.1 mm lateral and 1.3 mm anterior to lambda. The headframe was then affixed to the skull using C&B Metabond (Parkell). Once the Metabond was dry, the skull was rotated by 20° to make a circular craniotomy (5 mm diameter) over visual cortex. After removal of the skull, a durotomy was performed and a circular glass coverslip (5 mm diameter with additional 1 mm lip to affix to skull) was lowered into the craniotomy and affixed to the skull with Vetbond. The bottom of the coverslip was coated with a layer of polydimethylsiloxane (SYLGARD 184, Sigma-Aldrich) to reduce adhesion to the brain surface. A ring of Kwik-Cast was added along the perimeter of the coverslip and held in place by thin Metabond bridges between the coverslip and headplate well. Following surgery, mice were monitored for overall health and cranial window clarity over a 7-10 day recovery period.

Four mice were implanted with SHIELD implants^3^ instead of glass coverslips to permit recordings over four consecutive days.

### Intrinsic Signal Imaging

Intrinsic signal imaging (ISI) was used to measure the hemodynamic response of the cortex to visual stimulation across the entire cranial window^2^. ISI results were then used to delineate visual cortical area boundaries and guide probe insertions.

Mice were lightly anesthetized with 1-1.4% isoflurane and eye drops were applied (Lacri-Lube Lubricant Eye Ointment). Mice were headfixed for imaging normal to the cranial window. Before ISI, an image of the vasculature was taken under green light to register ISI data to the brain surface. Next the imaging plane was defocused and the hemodynamic response to a visual stimulus was imaged under red light. The visual stimulus was an alternating checkboard pattern (25° square size) superimposed on a 20° wide bar drifting across a mean luminance gray background. On each trial, the stimulus bar was swept across the four cardinal axes 10 times in each direction at a rate of 0.1 Hz.

A minimum of three trials were averaged to produce altitude and azimuth phase maps, calculated from the discrete Fourier transform of each pixel at the stimulus frequency. A “sign map” was produced from the phase maps by taking the sine of the angle between the altitude and azimuth map gradients. In the sign maps, each cortical visual area appears as a contiguous red or blue region (Figure 1a).

### Behavior Training

#### Water restriction and habituation

Throughout training mice were water-restricted to motivate learning and performance of the behavioral task. Mice had access to water only during behavioral training sessions or when provided by a technician on non-training days. Before water restriction, mice were weighed for three consecutive days to establish a baseline weight. During the first week of water restriction mice were habituated to daily handling and increasing durations of head fixation in the behavior enclosure over a five-day period. Mice were trained 5 days per week (Monday-Friday) and were allowed to earn unlimited water during the daily 1 hour sessions. Supplemental water was supplied in the home cage if the volume earned during the task fell below 1 mL or the body weight fell below 85% of the baseline weight. On non-training days mice were weighed and received enough water to reach 85% of their baseline weight (but never less than 1 mL).

#### Apparatus

Mice were trained in custom built behavior enclosures equipped with a 24” gamma-corrected LCD monitor (ASUS, #PA248Q). Mice were head-fixed on a behavior stage and allowed to run on a 6.5” running wheel. The center of the visual monitor was placed 15 cm from the right eye and visual stimuli were spherically warped to correct for distortion at the periphery of the monitor. Water rewards were delivered using a gravity-fed solenoid (NI Research, #161K011) to deliver a calibrated volume of water through a blunted 17g hypodermic needle (Hamilton) positioned approximately 2 mm away from the animal’s mouth using a custom 3-axis motorized stage.

#### Change Detection Task

This task has been previously described in detail^4,5^. Briefly, mice were trained in a go/no-go change detection task (schematized in Figure 1f). Mice were presented with a continuous stream of natural images (250 ms duration followed by 500 ms of isoluminant gray screen) and were trained to lick a reward spout when the identity of the stimulus changed. If mice responded correctly within a short, post-change response window (150-750 ms after stimulus change) a water reward was delivered. A ‘grace period’ of 3 seconds occurred after each change to allow for water consumption. During this period no additional changes could occur. At the start of each trial, the ‘change’ stimulus was selected. Given that there were eight total images, ∼7/8 of trials were ‘GO’ trials (the ‘change’ stimulus selected differed from the current stimulus) and 1/8 were ‘CATCH’ trials (the ‘change’ stimulus selected was the same as the current stimulus). A change time was selected from a geometric distribution ranging from 5 to 12 flashes. If the mouse licked prior to the stimulus change the trial restarted (‘aborted’). If a given trial was aborted 5 times, a new trial and change time were drawn. No other punishment was delivered for aberrant licking. Licks during the response window were scored as ‘hits’ on GO trials and ‘false alarms’ on CATCH trials. During electrophysiology experiments but not during prior training, approximately 5% of image stimuli were omitted, extending the gray screen between stimuli from 500 ms to 1.25 ms. The stimulus immediately preceding a change could not be omitted, and two consecutive stimuli were never omitted.

#### Automated Training

Mice were trained using an automated procedure that comprised six stages. On the first training day (Stage 0), mice were introduced to the rig and the lickspout during a 15 minute ‘open-loop’ session. Non-contingent 10 µL water rewards were delivered coincident with 90° changes in the orientation of a full-field static square-wave grating. All subsequent training stages were run in ‘closed-loop’ and required the mouse to lick for water reward as follows:

Stage 1: Mice were presented with static, full-field square wave gratings (oriented at 0° or 90°) and rewarded with 10 µL of water for licking when the orientation of the grating changed. Progression to the next stage required a d-prime of at least 2 for 2 of the last 3 sessions.

Stage 2: A 500 ms inter-stimulus gray screen was introduced between grating stimuli (250 ms duration; oriented at 0° or 90°). Progression criteria were identical to Stage 1.

Stage 3: Flashed gratings were replaced with natural images. Progression was triggered after 3 sessions.

Stage 4: Reward volume was reduced to 7 µL. Progression required a peak d-prime (calculated over a rolling window of 100 trials) greater than 1 for 3 consecutive sessions and at least 100 contingent (non-aborted) trials.

Stage 5: A 5 minute epilogue was appended at the end of the session, introducing mice to the gabor stimuli and full field flashes used to map receptive field properties. Reward volume was reduced to 5 µL. A five minute waiting period was added to the beginning of the session to acclimate mice to experiment preparation time during electrophysiology experiments. Mice were eligible to be handed off for electrophysiology when they achieved a mean reward number of 120 over the last 3 sessions in addition to the Stage 4 criteria.

After training, mice were habituated to the electrophysiology rigs for 5-10 days before experiments during which the water reward was further reduced (3-5 µL) to prevent satiation.

In a previous dataset from the Allen Institute using the same change detection task^5^, many mice relied at least partially on a timing strategy, licking more than expected by chance about 5 stimuli from the end of the last trial (i.e. the first eligible change image presentation)^6^. To discourage mice from suboptimal strategies, we added a strict reward threshold to the passing criterion for the final training stage. Applying the same dynamic logistic regression model used in Piet et al. (2024)^6^, we quantified the impact of the added criterion with a strategy index score, indicating whether the mouse relied more heavily on visual (positive score) or timing (negative score) cues. Indeed, we found that the mice in this dataset were significantly more biased towards a visual strategy than mice in the previous dataset (strategy index mean ± SEM: 9.5 ± 1.3 for this dataset, -5.1 ± 0.59 for Garrett et al. (2023)^5^; p=1.2e-23, Wilcoxon rank-sum test).

### Neuropixels Recordings

#### Preparation of the mouse for recording

After the completion of a successful ISI map, a custom insertion window was generated for each mouse. First, six insertion targets were manually drawn on the V1-aligned eccentricity map using a web-based annotation tool. Targets were positioned at the center of retinotopy for each area. The coordinates of each target were used to automatically generate the outlines of the insertion window, which was subsequently laser-cut out of 0.5 mm clear PETG plastic (Ponoko). When seated in the headframe well, the window facilitates access to the brain via holes over each of the six visual areas.

On the morning of the first recording day, the cranial coverslip was removed and replaced with an insertion window containing holes aligned to six cortical visual areas as previously described^2,7^. Briefly, on the morning of the first recording day, the mouse was anesthetized with isoflurane (3-5% induction and 1.5% maintenance) and the eyes were protected with ocular lubricant. Body temperature was maintained at 37.5°C. The Kwik-Cast surrounding the cranial window was removed and a silicone suction cup was attached to the window. If necessary, a bent syringe needle was used to free the coverslip from Vetbond, and the cranial window was lifted from the brain surface. The insertion window was then placed above the brain and affixed to the headframe well with Metabond. An agarose mixture was injected underneath the window and allowed to solidify. The mixture consisted of 0.4 g high EEO Agarose (Sigma-Aldrich), 0.42 g Certified Low-Melt Agarose (BioRad), and 20.5 ml ACSF (135.0 mM NaCl, 5.4 mM KCl, 1.0 mM MgCl_2_, 1.8 mM CaCl_2_, 5.0 mM HEPES). A layer of silicone oil was added over the holes in the insertion window to prevent the agarose from drying out. A plastic cap was screwed into the headframe well to protect the preparation from cage debris. After this procedure (∼30 minutes), mice were returned to their home cage and allowed to recover for at least 2 hours (typically 3-4) before the recording experiment.

After the first recording session, the probes were retracted, silicone oil was removed from the window surface, and the well was sealed with additional agarose added on top of the implant window. The protective cap was re-attached and the mouse was returned to its home cage. On the following day, the mouse was returned to the rig, the excess agarose was removed from the implant window and a fresh layer of silicone oil was applied before reinsertion of the probes for the second recording. Receptive fields generated from the first recording session for each visual area were used to adjust the probe insertion targets if necessary to maximize RF overlap with the center of the screen.

For mice with SHIELD implants, the mouse was briefly anesthetized on the Friday before the experimental week, and the SORTA-Clear plug protecting the implant was removed and replaced with Kwik-Cast. On the morning of each recording day, the Kwik-Cast plug was removed and the implant was covered with agarose and a thin layer of silicone oil. At the end of the recording, the agarose was removed and replaced with a fresh layer of Kwik-Cast.

#### Probes

All neural recordings were performed with Neuropixels 1.0 probes^8^. The electrodes closest to the tip were always used, providing a maximum of 3.84 mm of tissue coverage. All recordings were referenced to the large pad at the tip of the probe. Our goal was to insert six probes per experiment.

Each probe was coated in CM-DiI (1 mM in ethanol; Thermo Fisher, V22888) to visualize probe tracks after tissue clearing and ex vivo imaging. The probes were dipped in a well filled with the dye for approximately 1 minute.

#### Head fixation

The mouse was placed on the running wheel and fixed to the headframe clamp. The plastic cap was removed from the headframe well and an aluminum cone was lowered to prevent the mouse’s tail from striking the probes. An infrared dichroic mirror was placed in front of the right eye to allow the eye-tracking camera to operate without interference from the visual stimulus. A black curtain was then lowered over the front of the rig, placing the mouse in complete darkness except for the visual stimulus monitor.

#### Grounding

A 32 AWG silver wire (A-M Systems) was epoxied to the headframe before the initial headframe/cranial window surgery. This wire becomes electrically conductive with the brain surface after the application of the ACSF/agarose mixture beneath the insertion window. The wire was pre-soldered to a gold pin embedded in the headframe well, which mates with a second gold pin on the protective cone. The cone pin was soldered to 22 AWG hook-up wire (SparkFun Electronics), which was connected to both the behavior stage and the probe ground. Before the experiment, the brain-to-probe ground path was checked using a multimeter.

The reference connection on the Neuropixels probes was permanently soldered to ground using a silver wire, and the headstage grounds (which are contiguous with the Neuropixels probe grounds) were connected in parallel to the animal ground.

#### Probe Insertion

After the mouse was secured in the headframe, the cartridge holding the probes was lowered so the probe tips were approximately 2.5-mm above the brain surface. The probes were then manually lowered one by one to the brain surface until spikes were visible on the electrodes closest to the tip. After the probes penetrated the brain to a depth of around 100 μm, they were inserted automatically at a rate of 200 μm min^−1^ (total of 3.5 mm or less into the brain) to avoid damage caused by rapid insertion. After the probes reached their targets, they were allowed to settle for at least 10 minutes.

#### Data acquisition and synchronization

Neuropixels data was acquired at 30 kHz (spike band) and 2.5 kHz (LFP band) using the Open Ephys GUI. Gain settings of 500× and 250× were used for the spike band and LFP band, respectively. Probes were connected to a PXIe card inside a National Instruments chassis (Neuropixels 1.0). Raw neural data was streamed to a compressed format for archiving, which was extracted before analysis.

Videos of the eye, body and face were acquired at 60 Hz. The angular velocity of the running wheel was recorded at the time of each stimulus frame, at approximately 60 Hz. Synchronization signals for each frame were acquired by a dedicated computer with a National Instruments card acquiring digital inputs at 100 kHz, which was considered the master clock. A 32-bit digital ‘barcode’ was sent with an Arduino Uno (SparkFun DEV-11021) every 30 s to synchronize all devices with the neural data. Each Neuropixels probe has an independent sample rate between 29,999.90 Hz and 30,000.31 Hz, making it necessary to align the samples offline to achieve precise synchronization. The synchronization procedure used the first matching barcode between each probe and the master clock to determine the clock offset, and the last matching barcode to determine the clock scaling factor.

#### Lick Detection

Licks were detected with a capacitive lick sensor during training. However, this sensor was incompatible with Neuropixels recordings due to the electrical artifact generated during licking. To address this issue, a custom piezo lick sensor was designed for electrophysiology rigs consisting of a non-conductive PEEK needle (Hamilton) resting on a piezo microphone (TE Connectivity, 1007079-1). Licks were detected by the physical displacement of the spout during contact with the tongue. To add rigidity, the PEEK needle was reinforced either with a wooden applicator or by a stainless-steel sleeve (Hamilton) which stopped approximately a centimeter from the end of the needle. The analog microphone signal was digitized with a SAMD21 microprocessor and band-pass filtered between 3 and 35 Hz. The resulting signal was then passed through a software comparator to generate the final TTL lick signal.

#### Stimulus Monitor

Visual stimuli were generated using custom scripts based on PsychoPy^9^ and were displayed using an ASUS PA248Q LCD monitor, with 1,920 × 1,200 pixels (55.7 cm wide, 60 Hz refresh rate). Stimuli were presented monocularly, and the monitor was positioned 15 cm from the right eye of the mouse and spanned 120° × 95° of visual space before stimulus warping. Each monitor was gamma corrected and had a mean luminance of 50 cd m^−2^. To account for the close viewing angle of the mouse, a spherical warping was applied to all stimuli to ensure that the apparent size, speed and spatial frequency were constant across the monitor as seen from the mouse’s perspective.

#### Experimental design and visual stimuli

Every experiment followed the timeline diagrammed in Figure 1e. Briefly, mice encountered the following stimulus blocks (in order of appearance):

Active behavior block: the mouse performed the detection of change task described above. The images used during this block came from one of two image sets of 8 natural images (G or H). On one recording day, mice were exposed to the same image set they had seen during behavior training (‘Familiar’ sessions). On the other recording day, mice were shown a novel image set (‘Novel’ sessions). Two images were shared across these image sets (‘holdover’ images) and were thus familiar on both sessions. In total, 41 mice were trained on image set G, and shown G on the first recording day and H on the second; 10 mice were trained on H, and shown H on the first recording day and G on the second; 3 mice were trained on G and shown H on the first recording day and G on the second.

Images were presented for 250 ms, with a 500 ms gray screen between images. During recording sessions, images were omitted (resulting in a gray screen lasting 1.25 seconds) with a probability of 5%. However, the image immediately preceding the change image and the image immediately following an omission were never omitted, producing a true omission probability of ∼3%. At the end of this block, the lick spout was retracted for the remainder of the session.

Gabor stimuli: Gabor stimuli windowed to have a 20° diameter were presented in a 9x9 grid to map receptive fields. Grid positions were spaced by 10 degrees. Gabors had a temporal frequency of 4 Hz, a spatial frequency of 0.08 cycles/degree and were shown at 3 orientations (0, 45, 90 degrees). They were 250 ms in duration without gaps between stimuli. There were 15 trials for each condition (81 positions, 3 orientations).

Gray screen: Following the Gabor stimuli, there were 5 minutes of gray screen without stimuli to measure spontaneous activity.

Full-field flashes: Flashes were black or white at 80% contrast. They were 250 ms in duration with 2 seconds between flash starts. There were 75 trials for each condition (light and dark flashes).

Passive replay: The stimuli encountered by the mouse during the active behavior block were re-shown frame-for-frame. Note however that the lick spout was retracted for this block and the mouse was therefore unable to earn rewards.

#### Probe removal and cleaning

After each experiment, the probes were retracted from the brain at a rate of ∼1mm/s and the mouse was removed from head fixation and returned to its home cage. The probes were then immersed in a well of 1% Tergazyme for ∼12 hours.

#### Quality Control for Neuropixels recording sessions

Out of a total of 180 sessions, 27 were excluded from the release due to the following QC failures: white foam buildup on the edge of the eye partially covering the pupil (1), software failures compromising critical data streams (2), visual stimulus synchronizing failure (2), cortical bleeding or compromised brain health resulting in low unit activity and/or atypical visual responses (18), gap in data acquisition (2), discovery of purulent material over right hemisphere during ex-vivo imaging (2).

Of the remaining 153 sessions, 50 were released with a quality control flag and omitted from the analysis in this manuscript. We flagged two potential abnormalities as follows:

Abnormal histology: Ex-vivo imaging revealed potential tissue damage in 27 mice. Individual brains were flagged for 1) discoloration indicating large bleeds or hypoxic tissue, 2) swelling of the hippocampus, 3) tissue bruising, or 4) skull deformity compressing the right cortex (contralateral to window). All recording sessions were flagged for these mice, as acute events like large subcortical bleeds or diffuse bruising could not be confidently ascribed to the first or second recording day (49 sessions total).

Abnormal activity: We flagged 12 sessions as having potential epileptiform activity, characterized by a sharp increase in the firing rate across the probe (lasting 1-5 seconds) followed by a sustained suppression of activity (lasting several minutes) for at least one probe during the session. Of these 12 sessions, 11 were also flagged for abnormal histology.

### Optotagging

#### Optotagging Protocol

At the end of every experiment (regardless of genotype), an optotagging protocol was run during which the cortical surface was stimulated with blue light. In *Sst-IRES-Cre;Ai32* and *Vip-IRES-Cre;Ai32* mice, this protocol allowed us to identify putative Sst+ and Vip+ cortical interneurons by a dramatic increase in spiking activity time-locked to laser stimulation (and consequent ChR2 activation). We chose to focus on these two cell classes because they make up two of the three major non-overlapping populations of cortical interneurons^10^. The third major class (parvalbumin-expressing cells), can largely be distinguished by their narrow spike waveforms^11^. Blue light was delivered by a 473 nm laser (Laser Quantum, model Ciel or Cobolt model 06-MLD). The light source was coupled to a 400 μm diameter fiber optic cable (Thorlabs) or bifurcated fiber bundle (Thorlabs, BFYL4LF01), with the tip(s) positioned such that blue light illuminated the entire cranial window. Two types of stimuli at 3 different light levels were randomly interleaved: a 10 ms pulse, and a 1s raised cosine ramp. For the pulse stimulus, a 0.5 ms ramp was applied at the beginning and end of the pulse (Figure 5). Stimuli were presented at intervals of 1.5 s plus a uniformly distributed delay between 0 and 0.5 s.

#### Cell type label assignment

To assign putative cell type labels to units recorded in *Vip-IRES-Cre;Ai32* or *SST-IRES-Cre;Ai32* mice, we defined criteria based on the following metrics characterizing each unit’s response to the laser stimuli. Note that, for all analysis, we ignored the first 1.5 ms after laser onset and the last 1.5 ms before laser offset to avoid potential light artifacts in the recording, yielding 7 ms for analysis of the 10 ms laser pulse and 997 ms for the 1 second cosine ramp stimulus. We defined four criteria, requiring that optotagged neurons exhibit fast, reliable spiking to the laser pulse onset and sustained firing for the long cosine-ramp stimulus.

Z-scored response to the 10 ms pulse: The baseline spike rate was determined by randomly drawing 7 ms windows from the intertrial intervals (ignoring activity within 5 ms of laser onset and offset) and counting spikes for these bins. We required that the mean spike count during the high power 10 ms pulse stimulus be greater than the mean rate over these baseline windows plus two standard deviations.

Response latency for the 10 ms pulse: For each trial of the high power 10 ms pulse condition, we measured the time of the first spike after laser onset. We required that the median latency be less than 8 ms.

Response jitter for 10 ms pulse: We required that the median absolute deviation of first spike times as defined above be less than 2 ms.

Fraction of time with significantly elevated spike rate for the cosine ramp stimulus: Baseline windows (997 ms duration) were randomly drawn from intertrial intervals as described above and spikes were counted in 10 ms bins for each window. The response to the high power cosine stimulus was also binned in 10 ms bins. For each bin, a Mann-Whitney U test was performed to compare the spike counts from the baseline and stimulus windows. We required that at least 30% of the bins be significantly (p < 0.01) elevated from baseline. Note that this is a conservative criterion and may exclude ChR2-expressing neurons with only a transient response to the laser.

In addition to these cell type labels, we defined regular spiking (RS) units as those that did not pass the above criteria and had a trough-to-peak duration of greater than 0.4 ms for the mean spike waveform at the peak channel (the channel with the greatest difference between the peak and trough). Fast spiking (FS) units were required to have a peak-to-trough duration of less than 0.4 ms.

### Ex Vivo Imaging

#### Tissue Clearing

Mice were anesthetized (5% isoflurane for induction), then perfused with 4% paraformaldehyde (PFA). The brains were preserved in 4% PFA, rinsed with PBS the next morning, and stored at 4°C in PBS. A modified iDISCO method was then used to clear the brains^12^. On the first day, the brains were immersed in increasing concentrations of methanol (20, 40 and 60%) for an hour each, then overnight in 80% methanol. On the second day, they were dipped in 100% methanol (twice for one hour) and then into a mixture of 1/3 methanol and 2/3 dichloromethane overnight. On the third day, the brains were moved from pure dichloromethane (2 × 20 min) to pure dibenzyl ether, where they remained for 2-3 days until clearing was complete.

#### Optical projection tomography

Whole-brain 3D imaging was accomplished with optical projection tomography (OPT), as described previously^2^. The OPT instrument consisted of collimated light sources for transmitted illumination (on-axis white LED, Thorlabs MNWHL4 with Thorlabs SM2F32-A lens and Thorlabs DG20-600 diffuser) or fluorescence excitation (off-axis Thorlabs M530L3, with Thorlabs ACL2520U-DG6-A lens and Chroma ET535/70m-2P diffuser), a 0.5× telecentric lens (Edmund Optics 62-932) with emission filter (575 nm LP, Edmund Optics 64-635), and a camera (IDS UI-3280CP). The specimen was mounted on a rotating magnetic chuck attached to a stepper motor, which positioned it on the optical axis within a glass cuvette filled with dibenzyl ether. The stepper motor and illumination triggering were controlled with an Arduino Uno (SparkFun DEV-11021) and custom shield including a Big Easy Driver (SparkFun ROB-12859). Instrument communication and image capture was accomplished with MicroManager.

A series of 400 images were captured with transmitted LED illumination. The specimen was rotated 0.9° between images (thus making one full rotation through the image series). The same procedure was repeated with the fluorescence excitation LED. Each channel was stored as a separate OME-TIFF dataset before extracting individual planes and metadata required for reconstruction using a custom Python script.

Isotropic 3D volumes were reconstructed from these projection images using NRecon (Bruker). The rotation axis offset and region-of-interest bounds were set for each image series pair using the transmitted channel dataset, then the same values applied to the fluorescence channel dataset. A smoothing level of 3 using a Gaussian kernel was applied to all images. Reconstructions were exported as single-plane 16-bit TIFF images taken along the rotation axis with final voxel size of 7.9 μm per side.

#### Registering probes to the common coordinate framework

Reconstructed brains were registered to the Allen Institute Common Coordinate Framework (CCFv3) as previously described^2^. Reconstructed volumes were downsampled to 10 μm per voxel and roughly aligned to the CCFv3 template brain using an affine transform. The volume was then cropped to a size of 1,023 × 1,024 × 1,024 and converted to Drishti format (https://github.com/nci/drishti). Next, 6–54 registration points were marked in up to 14 coronal slices of the individual brain by comparing to the CCFv3 template brain. Fluorescent probe tracks were manually labelled in coronal slices of the individual brain, and the best-fit line was found using singular value decomposition. The registration points were used to define a 3D nonlinear transform (VTK thinPlateSplineTransform), which was used to translate each point along the probe track into the CCFv3 coordinate space. Each CCFv3 coordinate corresponds to a unique brain region, identified by its structure acronym (for example, CA3, LP, VISp, etc.). A list of CCFv3 structure acronyms along each track was compared to the physiological features measured by each probe (for example, unit density, LFP theta power, visual responsiveness). The locations of major structural boundaries were manually identified to align the CCFv3 labels with the physiology data. After the manual alignment procedure, each recording channel (and its associated units) was assigned to a unique CCFv3 structure. White matter structures were not included; any units mapped to a white matter structure inherited the grey matter structure label that was immediately ventral along the probe axis.

Because two recordings were made in most animals (maximum of 12 insertions in each brain), information about the relative position of each probe on the first and second insertions was required to assign an annotated probe track to its appropriate recording session. Two data sources were consulted to make this judgement: 1) the operator’s record of where each probe was inserted relative to surface vasculature landmarks for each recording day (annotated as described in *Identi;cation of cortical visual area targets*), and 2) the record of the movement of each probe’s manipulator. Together, these records were used to estimate the position of a given probe on the first recording day relative to the second and thus assign annotated probe tracks to their corresponding session.

#### Identification of cortical visual area targets

To confirm the identity of the cortical visual areas, images of the probes taken during the experiment were compared to images of the brain surface vasculature taken during the ISI session. Key points were selected along the vasculature on both images, and a perspective transform (OpenCV) was performed to warp the insertion image to the retinotopic map. The probe entry points were then manually annotated, and an area was assigned to each probe based on the ISI-derived visual area boundaries. Penetration points that could not be unambiguously associated with a particular visual area were classified as ‘VIS’. If the cortical area label obtained via CCFv3 registration did not match the area identified in the insertion image overlay, the insertion image overlay took precedence.

#### Cortical layer labels

CCFv3 coordinates were used as indices into the template volume to extract layer labels for each cortical unit (L1, L2/3, L4, L5, or L6). The relative thickness of each layer, which can vary both within and across areas, is based on the average of the 1,675 individual brains used to create the template volume.

#### Probe annotation quality control

For 41 out of 905 insertions, probe tracks could not be confidently annotated in the imaging volume. For these tracks, the beginning and end of cortex were identified using electrophysiological hallmarks (e.g. unit density, LFP power, visual responsiveness), and cortical units were assigned to regions based on ISI as described above. All subcortical units for these probes were assigned to ‘grey’.

### Spike Sorting

To preserve consistency with the Allen Institute Visual Coding Neuropixels data release, all data processing and spike-sorting was performed according to the pipeline described previously (https://github.com/AllenInstitute/ecephys_spike_sorting)^2^.

#### Data pre-processing and spike-sorting

Before spike-sorting, the spike-band data passed through four steps: offset removal, median subtraction, filtering and whitening. First, the median value of each channel was subtracted to center the signals around zero. Next, the median across simultaneously sampled channels was subtracted to remove common-mode noise. For each sample, the median value of channels *n*:24:384, where *n* = [1,2,3,…,24], was calculated, and this value was subtracted from the same set of channels. The original data were overwritten with the median-subtracted version, with the median value of each block of 16 channels saved separately, to enable reconstruction of the original signal if necessary. The median-subtracted data file is sent to the Kilosort2 MATLAB package (https://github.com/mouseland/Kilosort2, commit 2fba667359dbddbb0e52e67fa848f197e44cf5ef)^13^, which applies a 150-Hz high-pass filter, followed by whitening in blocks of 32 channels. The filtered, whitened data are saved to a separate file for the spike-sorting step.

#### Spike-sorting quality control

Units were automatically classified as noise based on three criteria: waveform spread either restricted to a single channel or extending over more than 25 channels, waveform shape (no peak and trough, based on wavelet decomposition), or multiple spatial peaks (waveforms are non-localized along the probe axis). A suite of waveform metrics was then calculated to further refine non-noise units. The following metrics were used to filter units for analysis in this study:

Presence ratio: The session was divided into 100 equal-sized blocks; the presence ratio is defined as the fraction of blocks that include one or more spikes from a particular unit. Units with a low presence ratio are likely to have drifted out of the recording or could not be tracked by Kilosort2 for the duration of the experiment. Units were required to have a presence ratio greater than 0.9.

ISI (interspike intervals) violation ratio: This metric searches for refractory period violations that indicate a unit contains spikes from multiple neurons. It is defined as the ratio of the rate of contaminating spikes to the overall spike rate for a given unit and is calculated by counting the number of violations of less than

1.5 ms, dividing by the amount of time for potential violations surrounding each spike, and normalizing by the overall spike rate. It is always positive (or 0), but has no upper bound. Units were required to have an ISI violation ratio less than 0.5.

Amplitude cutoff: This metric provides an approximation of a unit’s false negative rate. First, a histogram of spike amplitudes is created, and the height of the histogram at the minimum amplitude is extracted. The percentage of spikes above the equivalent amplitude on the opposite side of the histogram peak is then calculated. If the minimum amplitude is equivalent to the histogram peak, the amplitude cutoff is set to 0.5 (indicating a high likelihood that more than 50% of spikes are missing). This metric assumes a symmetrical distribution of amplitudes and no drift, so it will not necessarily reflect the true false negative rate. Units were required to have an amplitude cutoff of less than 0.1.

### VISp silencing experiment

To silence VISp activity, a multimode optic fiber (250 µm diameter, ThorLabs) was placed above the ISI-defined retinotopic center of V1 in VGAT-ChR2 mice expressing channelrhodopsin in cortical inhibitory neurons. The fiber was connected to a 488 nm blue laser calibrated to deliver ∼2.5 mW at the fiber tip. Light was delivered on 30% of trials, turning on at one of four intervals from stimulus change onset: -12 ms (range: -13, -11), 39 ms (37, 39), 72 ms (70, 72) and 105 (104, 106). One mouse licked reliably for the laser onset and was excluded from analysis.

### Data processing

#### Running wheel data

Running speed was monitored using an analog encoder mounted at the center of a 6.5” diameter running wheel. To correct for noise in the raw voltage signal, outlier values (more than 10 standard deviations from the mean) were removed and the final speed trace was lowpass filtered with a 10 Hz zero-phase Butterworth filter (scipy.signal.filtfilt^14^).

#### Video data

A standardized pipeline was built for fitting ellipses to the pupil, eye (visible perimeter of the eyeball), and corneal reflection of the right eye, based on points tracked using the open source software DeepLabCut (https://github.com/DeepLabCut/DeepLabCut) as previously described^5^. We used DeepLabCut, initialized with a pre-trained ResNet 50 deep residual network, to track (up to) 12 points along the perimeters of the eye, pupil, and corneal reflection. Ellipses were then fit to the tracking points. Validation against hand-annotated ‘ground truth’ frames confirmed that a single ‘universal’ model, trained on a broad selection of data samples, robustly generalized on held-out data across different physiology rigs and individual animals.

It was assumed that the pupil is round but when viewed obliquely appears as an ellipse, the major axis of which reflects the pupil diameter. Thus, the area of the pupil is calculated on every frame as the area of a circle with a diameter defined by the longest axis of the ellipse fit for that frame. The area of the ellipses for both the corneal reflection and eye ellipse fits are calculated using the standard formula for the area of an ellipse. On frames where the animal is blinking, tracked points may be missing or the confidence of tracked points may be low, and ellipse fits either fail completely or fit erroneously. To avoid including these erroneous fits in analysis, an algorithm attempts to identify these ‘likely blink’ frames. Likely blinks are identified as frames where either the eye or pupil fit is missing, or where the z-scored value of the eye or pupil area exceeds 3. In addition, two frames before and after every likely blink identified by the above methods are also labeled as likely blinks. This is to avoid the possibility of analyzing erroneous fits caused by a partially opened eye. Note that this algorithm may flag not only genuine blink frames but also frames where the DeepLabCut algorithm simply failed to identify a reasonable fit.

### Data analysis

For all analyses, only non-noise units from sessions without QC flags and which passed the presence ratio, ISI violation ratio and amplitude cutoff criteria were included.

#### Receptive Fields

Receptive fields (Figure 1) were calculated for each unit by taking the mean spike rate in the 250 ms following the presentation of a gabor patch in each of 81 locations (arranged in a 9×9 grid; see *Experimental design and visual stimuli* above). For each session, receptive fields for units in each visual area were averaged together to create a population receptive field. A 2D gaussian was then fit to this population receptive field and the peak position was taken as the receptive field center.

#### Hit rate, false alarm rate and d-prime

To calculate behavioral response rates, trials were first filtered for engagement. Engaged trials were defined as those for which the reward rate was at least 2 rewards per minute over the preceding 25 trials. Hit rates were then calculated as the fraction of engaged ‘Go’ trials for which the mouse licked within the response window after the change stimulus. False alarm rates were defined as the fraction of engaged ‘Catch’ trials for which the mouse responded after a ‘sham change’ stimulus. To properly compare hit and false alarm rates, sham changes were drawn from the same geometric distribution as change stimuli. For the analysis in Figure 1h, trials were aggregated across the two recording days before analysis. For one mouse, response rates were too low to reliably estimate the false alarm rate and the data from this animal were excluded. D-prime was calculated as the difference between the inverse of the cumulative distribution function (scipy.stats.norm.ppf) at the hit and false alarm rates.

#### Regression Model

The regression model was adapted from that used in Garrett et al. (2023)^5^ to be compatible with electrophysiology data. Briefly, we fit neural activity with a linear regression model featuring time-dependent kernels. Each model feature was a vector *f*_*i*_(*t*), which was convolved with a learned kernel *k*_*i*_(*t*) to produce the predicted model component for that feature. Model components were then summed together to produce the full model response. For continuous features (running speed and pupil diameter) the feature vectors were timeseries. For discrete features, *f*_*i*_(*t*) was 1 on timesteps where that feature occurred and 0 elsewhere. The complete list of features and their associated kernel widths is as follows:

Hits: 1 at stimulus onset for change stimuli during ‘hit’ trials and 0 elsewhere; 1.5 second kernel width.

Misses: 1 at stimulus onset for change stimuli during ‘miss’ trials and 0 elsewhere; 1.5 second kernel width.

Omissions: 1 at onset of omitted stimuli (i.e. when the stimulus would have otherwise occurred) and 0 elsewhere; 1.5 second kernel width.

Running: continuous time series of running wheel data; 2 second kernel width.

Pupil: continuous time series of pupil diameter; 2 second kernel width.

Licks: 1 for time bins during which a lick was registered and 0 elsewhere; 2 second kernel width

Images (one for each unique image, totaling 8): 1 at stimulus onset for all presentations of the respective image and 0 elsewhere; 0.75 second kernel width.

The model thus took the form *y* = *Wx*, where y is the 1d time series for a given unit (spiking activity binned in 25 ms bins), x is a vector of all kernel weights concatenated together, and W is a Toeplitz matrix with diagonal bands that map kernel weights onto time-shifted features. To solve for the kernel weights, we added an L2 ridge regression penalty to the standard solution for ordinary least squares regression, resulting in the following solution:

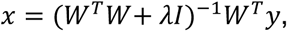

where λ is the L2 penalty. The model was fit with 5-fold cross-validation: we split each session into 50 intervals and randomly assigned 10 intervals to each cross-validation fold (each fold was intermingled in time). The hyper-parameter λ was selected for each session as the value (ranging from 0 to 500) that resulted in the best test-set performance across all units in that session. The training/test sets for hyper-parameter selection were different than those used for fitting the model for analysis.

To evaluate the unique contribution of each kernel to a given unit’s activity, we fit a series of reduced models where individual kernels or groups of kernels (components) were removed from the model and the model was refit. We then computed a dropout score, which measures the absolute contribution of a given feature *i* to the variance explained for each unit: *VE*_*full*_ − *VE*_*reduced*,*i*_.

#### Functional Clustering

Neurons were grouped into 12 clusters based on their linear regression kernel weights (see *Regression Model* section). The neurons with total explained variance less than 0.05 were automatically assigned to a “non-coding” cluster. We clustered the remaining units based on the following kernels: the preferred image for each unit, omissions, hits, misses, running, pupil diameter, and licks. The feature matrix for clustering was generated by concatenating the kernel weights for all responsive units (*N neuron x* 443 *weights*). An iterative algorithm consisting of an outer loop and an inner loop was then used for the clustering process. Briefly, for each step of the outer loop, the inner loop iterated through all existing clusters, attempting to split each one. These candidate splits were then evaluated with a silhouette score and the best was chosen, producing N+1 clusters for the next iteration of the outer loop.

In more detail, the inner loop attempted to split the *i*th cluster as follows:

1. Uniform Manifold Approximation and Projection (UMAP) was initialized with randomly selected parameters to reduce the feature matrix to 3 dimensions, resulting in an *N*_*i*_ *x* 3 matrix, where *N*_*i*_ is the number of units in the *i*th cluster.
2. A random sample of 5000 units was selected (if *N*_*i*_ was less than 5000, all units were used)
3. Spectral clustering (scikit-learn^15^) with random seed initialization was used to divide the sample units into two clusters.
4. A random forest classifier (scikit-learn) was trained on the output labels of the sample units from step (4) to label the remaining units.
5. Steps 1-4 were repeated 21 times. The cluster assignments for all 21 iterations were saved for each unit. The pattern of cluster assignments for a given unit can be considered a 21-digit binary code. The two most common codes across neurons defined the two most consistent clusters over the 21 clustering iterations. Each neuron was assigned to the cluster (‘code’) with which it shared the most matching bits (Hamming similarity).

The outer loop was terminated based on a combination of the silhouette score and visual inspection of the clusters.

To assess the robustness of the clusters, we ran a bootstrapped classifier on the same feature matrix used for clustering. For each classification run (100 total iterations), we used UMAP to reduce the dimensionality of the feature matrix to *N*_*units*_ *x* 3. A Random Forest classifier (scikit-learn RandomForestClassifier) was then trained on the cluster labels of the training set (20% of the units, randomly selected for each iteration) and subsequently used to predict labels for the test set (remaining 80% of units). Finally, a confusion matrix was constructed from the test set predictions by pooling across all 100 iterations (Figure S5).

#### Peri-stimulus/event histograms

Unit spiking activity was first collected in a response tensor for each session as follows: for every stimulus in the behavioral task, spikes during the interval from stimulus onset to the next stimulus onset (750 ms) were binned in 1 ms bins and binarized, resulting in a Boolean tensor of shape [number of units × number of stimuli × 750]. Trials of the desired type were then selected from this tensor and averaged for each unit. Unit responses were further filtered by area, cell type, layer and/or cluster as appropriate and aggregated across sessions before averaging. Unless otherwise noted, unit responses were smoothed with a symmetric exponential filter (tau = 3 ms) before averaging across units.

For the lick-triggered PETHs in Figure 2, unit activity was aligned to lick times during non-change stimuli. Shuffled lick responses were computed to identify what fraction of lick-triggered activity might be attributed to task events (e.g. visual stimuli) that correlated with licking. To create shuffled lick responses, lick times relative to stimulus onset were shuffled and the lick triggered average was recomputed.

For run-triggered peri-event histograms (PETHs) in Figure 2, the above tensor was not used. Rather, acceleration and deceleration events were identified from the running wheel data, and the spiking activity for each unit was binned in 1 ms bins and aligned to these events and averaged. Acceleration events were identified as follows: 1 second windows of running data were extracted for which the mean speed in the first half was less than 1 cm/s and the mean speed for the second half was greater than 5 cm/s. For each window, the acceleration event time was then defined as the time of maximum difference between adjacent time points (if the speed at this point was less than 1 cm/s) or the last point for which the running speed was less than 1 cm/s between the time of the minimum speed and the time of maximum difference. Deceleration events were similarly defined but now for windows for which the speed reduced from greater than 5 cm/s in the first half to less than 1 cm/s in the second half. Within these windows, the deceleration event time was defined as the point of maximum deceleration (minimum speed difference at adjacent time points) after the peak speed was reached. Unit responses were smoothed with a symmetrical exponential filter (tau = 3 ms) before averaging. These PETHs were calculated during the passive viewing epoch to avoid the correlation between running and licking present during active behavior.

Unless otherwise noted, PSTHs were baseline subtracted: the mean activity during the 50 ms before stimulus onset was subtracted from the stimulus response for each unit before averaging across units. PSTHs were always computed using engaged behavior trials.

The PSTH for visual cortex units in Figure 3 was computed for change stimuli during hit trials. Units in all six visual cortical areas were combined.

PSTHs in Figure 4 were computed for change, pre-change and passive change stimuli as follows: change stimuli were taken from hit trials; pre-change stimuli were defined as the stimuli immediately preceding change stimuli for hit trials. Change and pre-change stimulus responses were computed for each image separately and then averaged across images for each unit. Passive change stimuli were calculated for the same change stimuli but during passive replay.

PSTHs to familiar and novel stimuli in Figure 5 were computed for change stimuli from hit trials. Only non-holdover images (i.e. the 6 images unique to each image set) were used to compute both familiar and novel image responses. The PSTHs for familiar and novel stimuli in Figure 6 were computed using non-change stimuli, defined as stimuli during engaged behavior that 1) occurred at least 5 presentations since the last change (and were therefore outside the reward consumption window), 2) were not omitted, 3) did not immediately follow an omitted stimulus, and 4) occurred at least two stimulus presentations from the last lick response.

#### Image, change and lick decoding

Decoding from sensory and action clusters in Figure 3 was performed on pseudopopulations of neurons aggregated across sessions as follows:

For each session, neurons of the relevant cluster and region were identified, and stimulus presentations meeting the decoding category criteria were selected as illustrated in Figure 3b. Briefly, for image decoding, non-change presentations of each of the eight images were selected for which the mouse did not lick; for change decoding, non-change stimuli for which the mouse licked were compared to change stimuli for which the mouse licked; and for lick decoding, non-change stimuli for which the mouse did not lick were compared with non-change stimuli for which the mouse did lick. These stimuli were selected to isolate the axis of interest for each decoding analysis (image identity, change vs non-change, and lick vs no lick). All stimuli were taken from periods of engaged behavior (defined as trials during epochs for which the reward rate was >= 2 rewards over the preceding minute).

These unit and stimulus presentation ids were then used to index the session units × stimuli × 750 ms tensor described above in *Peri-stimulus/event histograms*. Activity was binned in 10 ms bins and aggregated in a list of tensors across sessions. The qualifying unit ids for each session were concatenated across sessions, and a pseudopopulation of 100 units was selected randomly from this list with replacement. For each unit in the pseudopopulation, the corresponding session tensor was loaded and that unit’s activity was extracted for all trials in each of the relevant decoding categories (for example, change and non-change trials for change decoding). For each trial category, trial activity for that unit was randomly assigned to train and test splits (50/50 split). Units from the same session were constrained to have the same train and test splits. Pseudotrials were then constructed across all units in the pseudopopulation by randomly selecting, for each unit, 100 trials for each trial category with replacement. Training data was constructed from each unit’s training split and testing data from each unit’s test split.

Once train and test pseudotrials were constructed, individual decoders were trained on progressively longer intervals of activity from stimulus onset (for example, the first decoder saw only the first 10 ms bin after stimulus onset, but the fifth decoder saw 5 bins capturing the first 50 ms of activity). Decoders were linear support vector classification models (scikit-learn’s LinearSVC with C = 1.0 and class_weight = ‘balanced’). This entire process was repeated for 100 randomly selected pseudopopulations, and the balanced accuracy for test data was averaged across runs to quantify decoding performance.

To test whether the time lag between LP and VISp change decoding might reflect different sampling of units from the two regions across sessions (Figure S8d,e), we subsampled sensory cluster units to match the number taken from each region for each session as follows: 1) sessions with at least one unit from each region were identified; 2) for each of these sessions, units were counted for each region and the minimum count across regions was identified; 3) this minimum count was randomly subsampled without replacement from the units in each region (i.e. all units were taken from the region with a lower unit count, and the same number were subsampled from the region with a greater unit count). After this matching process, units were pooled across sessions and decoding was run as described above.

#### Facemap lick decoding

To decode licking activity from video data (Figure 3), the side view video for each session was cropped to capture the face. The resulting cropped video was then processed with Facemap^13,16^ to extract the top 500 motion SVDs. Lick and non-lick trials were selected and a linear SVC decoder was trained on progressively longer video intervals from stimulus onset as described above. Decoders were trained with five-fold cross validation and performance on test data was averaged across sessions.

#### Image decoding for familiar and novel sessions

To compare the time course of image decoding during familiar and novel sessions, populations of 40 units were subsampled for each session and region without replacement. The number of sampling iterations was selected such that there was a 99% chance of sampling each unit at least once (sessions with less than 40 units for a given region were excluded). For each subsample, unit activity was binned in 10 ms bins and linear SVC decoders were trained with five-fold cross-validation to predict image identity from progressively longer time windows as described above. Recall was calculated for each image separately (number of times an image was correctly predicted divided by the total number of times it was presented), and the median was taken across subsamples. Recall was averaged across images (excluding hold-over images) to produce decoding time courses for each session.

#### Change, state and novelty modulation indices

Change and state modulation indices for each unit were calculated on the mean activity during the decision window (20-100 ms after stimulus onset) after baseline subtraction (baseline was defined as the 50 ms preceding stimulus onset). These indices compare activity across two conditions and were computed as follows:

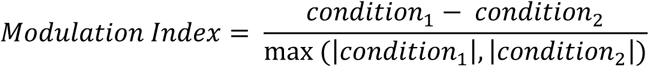

For change modulation, *condition*_1_ represents the mean firing rate during the decision window during change stimuli, and *condition*_2_ represents the mean firing rate during the decision window during pre-change stimuli. Both change and pre-change stimuli were taken from hit trials.

For state modulation, *condition*_1_ represents the response to change stimuli during active behavior, and *condition*_2_ represents the response to change stimuli during passive replay.

For each unit, the mean response was calculated for each image separately before averaging across images.

For novelty modulation, the index could not be calculated for individual units as different populations were recorded on the two recording days. Instead, units were filtered to meet the relevant criteria (e.g. region, cortical layer or cell-type) and separated according to whether they were recorded during the familiar session or the novel session. For each group, the mean response during the decision window was calculated for every unit as above and aggregated into a list. The familiar and novel responses were then resampled 1000 times with replacement. For each iteration, *condition*_1_ was defined as the mean response across the sampled novel units, and *condition*_2_ was defined as the mean response across the sampled familiar units. The final modulation index was then taken as the median value across iterations.

To examine the effect of running speed on state modulation (Figure S10), trials were subsampled to match the running speed distribution across active and passive epochs for each session before calculating unit-wise PSTHs and modulation indices as follows: 1) the mean running speed during a 200 ms window before each stimulus was binned in 5 cm/s bins separately for active and passive epochs; 2) for each bin, the minimum trial count was determined across the two epochs; 3) this minimum trial count was then subsampled without replacement from the trials in the corresponding bin for each epoch. Repeating this procedure for each bin produced identical running distributions for active and passive trials (Figure S10a).

#### Visual Response and novelty modulation latencies

To determine the visual and novelty modulation latencies for a given region, each unit’s mean response to non-change stimuli (excluding holdover images) and omitted stimuli were smoothed with a symmetrical exponential kernel (tau = 3 ms) and baseline subtracted. Units recorded during familiar and novel sessions were separated and a bootstrapped estimation of latency was calculated as follows: for each iteration, familiar image responses, novel image responses, familiar-session omission responses and novel-session omission responses were resampled across units with replacement. D-prime values were then calculated at each time point as follows: for visual latency, corresponding image and omission response distributions were compared; for novelty modulation latency, familiar and novel image response distributions were compared. Each comparison thus yielded a 250 element vector of d-prime values for the 250 ms stimulus window. The time of the maximum d-prime for this vector was identified, and the latency was defined as the first point working backwards from the maximum at which d-prime was <= 0.1, a threshold found empirically to identify the foot of the population visual response. If the maximum d-prime was less than 0.1, the latency was set to NaN for that iteration. For the final visual latency, familiar and novel visual latencies were averaged.

#### Change detection strategy analysis

To determine whether mouse behavior was better approximated by an ‘adaptation’ or ‘image-comparison’ strategy, the stimulus-wise responses of all units in a region of interest for a given session were averaged across the decision window (20-100 ms after stimulus onset), resulting in a response vector of length *N*_*units*_ for each stimulus presentation. The resulting response vectors were then used to train two decoding strategies as follows:

Adaptation decoder: A linear decoder (scikit-learn LinearSVC, with class_weight = ‘balanced’) was trained to classify stimuli as change or non-change. 5-fold cross-validation was used to assess the balanced accuracy of the decoder on held-out data.

Image comparison decoders: For each image, a one-vs-all decoder was trained with 5-fold cross-validation (scikit-learn LinearSVC, with class_weight = ‘balanced’) to differentiate stimulus responses to the selected ‘template’ image from responses to any of the other seven images (8 total decoders corresponding to 8 unique template images). For each stimulus presentation, the test-fold predictions of all 8 decoders were saved. The image-identity of the previous stimulus was then used to choose the appropriate decoder’s prediction and thus predict whether the current stimulus was the same as or different from the previous stimulus.

Applying these decoding strategies produced two vectors of length *N*_*stim presentations*_ for each session and region (depicted in Figure 7a). One vector stored the adaptation decoder confidence (scikit-learn’s decision function) that any given stimulus was a change stimulus. The other stored the image comparison decoder’s confidence that any given stimulus was the same as the previous stimulus.

These decoder confidence vectors were then used to predict two binary labels for each stimulus using logistic regression: *is_change* (whether a particular stimulus was a change stimulus, Figure 7b), and *is_lick* (whether the mouse licked for a particular stimulus, Figure S10d). For lick predictions, only non-change stimuli were considered, as model performance on these error trials was not confounded by there also being a stimulus change.

#### Other statistical methods

Unless otherwise stated, comparisons for a given metric across regions, cortical layers, cell-types or clusters were made using the Wilcoxon rank-sum statistic with each unit considered an independent sample (scipy.stats.ranksum). To correct for multiple comparisons, the Benjamini–Hochberg false discovery rate correction was applied (statsmodels.stats.multitest.multipletests with method ‘fdr_bh’).

