## Supplemental Figures for "Map of spiking activity underlying change detection in the mouse visual system"

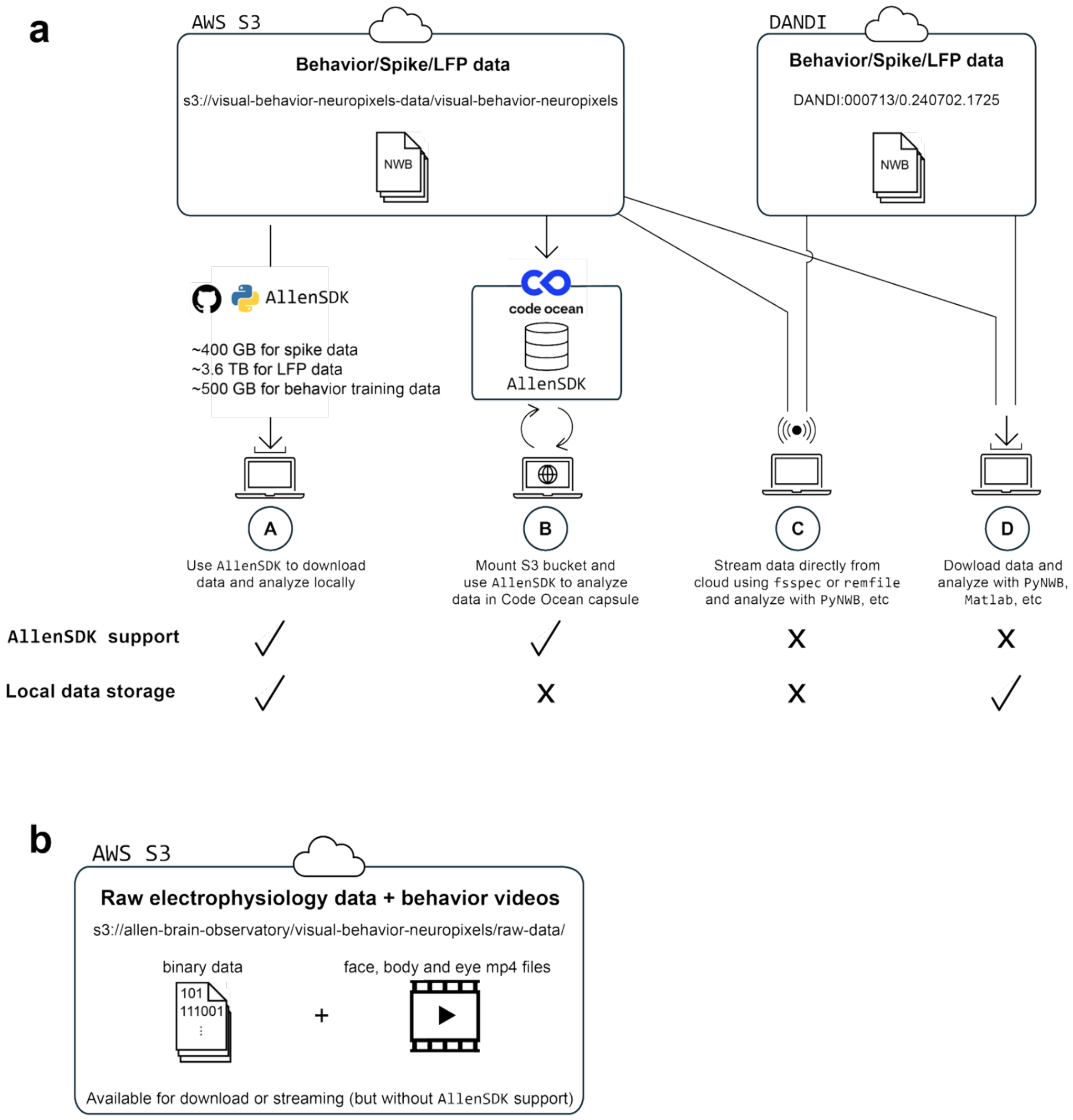


**Figure S1: Accessing the Allen Institute's Visual Behavior Neuropixels dataset. a)** Diagram depicting four different ways users might access this dataset. NWB files containing behavior, spike and LFP data are all stored in a public AWS S3 bucket as well as in the DANDI archive. Users can access this dataset in a number of ways, including: (A) Using the AllenSDK to download NWB files to local storage; (B) mounting the S3 bucket in Code Ocean to analyze on cloud compute resources, (C) using tools like fsspec or remfile to stream data directly from the cloud to be analyzed with PyNWB (or other packages capable of reading HDF5 files); or (D) downloading data to local storage for analysis without the AllenSDK. **b)** In addition to the NWB files above, raw electrophysiology data as well as behavior videos of the eye, face and body are available in a separate S3 bucket. The AllenSDK does not manage download or analysis of these data.

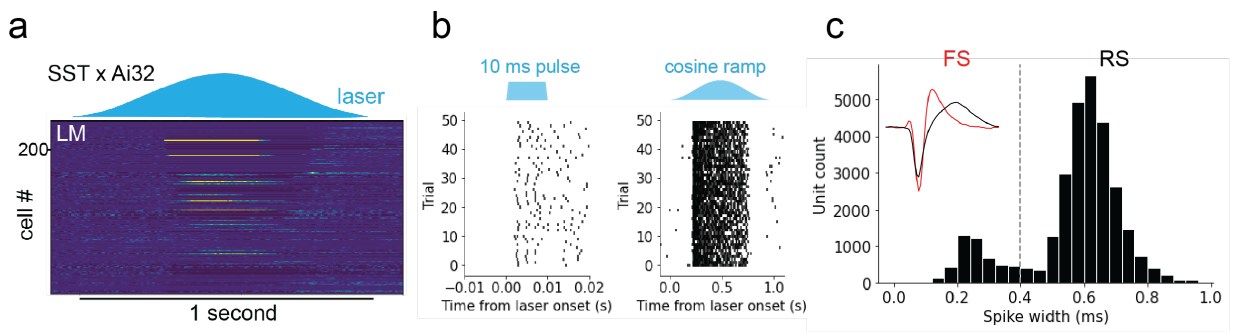


d

| cell type | Criteria | Count |
|---|---|---|
| RS | Not optotagged; spike duration > 0.4 ms | 25521 |
| FS | Not optotagged; spike duration < 0.4 ms | 4507 |
| SST | Optotagged in SST-cre x Ai32 mouse | 434 |
| VIP | Optotagged in VIP-cre x Ai32 mouse | 27 |

e

```
units['SST'] =  (units['genotype'].str.contains('Sst')) & \
                (units['pulse_high_mean_evoked_rate_zscored']>2) & \
                (units['pulse_high_first_spike_latency']<0.008) & \
                (units['pulse_high_first_spike_jitter']<0.002) & \
                (units['raised_cosine_high_fraction_time_responsive']>0.3)


units['VIP'] =  (units['genotype'].str.contains('Vip')) & \
                (units['pulse_high_mean_evoked_rate_zscored']>2) & \
                (units['pulse_high_first_spike_latency']<0.008) & \
                (units['pulse_high_first_spike_jitter']<0.002) & \
                (units['raised_cosine_high_fraction_time_responsive']>0.3)

units['RS'] =   (units['waveform_duration']>0.4)&(~units['SST'])&(~units['VIP'])
units['FS'] =   (units['waveform_duration']<0.4)&(~units['SST'])&(~units['VIP'])
```

**Figure S2: Identifying cortical cell types. a)** Heatmap showing mean response to cosine-ramp laser stimulus in SST-Cre x Ai32 mouse. Units (rows) are ordered by depth. Note several units with sustained response to laser (yellow). **b)** Raster for one example optotagged SST unit aligned to short pulse (left) and cosine ramp (right) laser stimuli. **c)** Histogram of peak-to-trough waveform duration for cortical units. Units with a duration of less than 0.4 ms were identified as FS. **d)** Table describing criteria and unit counts for each cortical cell type. **e)** Code snippet to demonstrate implementation of cell label criteria. Units were classified as optotagged if 1) their mean laser evoked firing rate was greater than two standard deviations above baseline for the short pulse at highest power, 2) the latency of their response to the pulse stimulus was less than 8 ms, 3) the jitter in their first spike during the pulse stimulus was less than 2 ms and 4) their firing rate was significantly elevated above baseline for at least 30% of the cosine ramp stimulus.

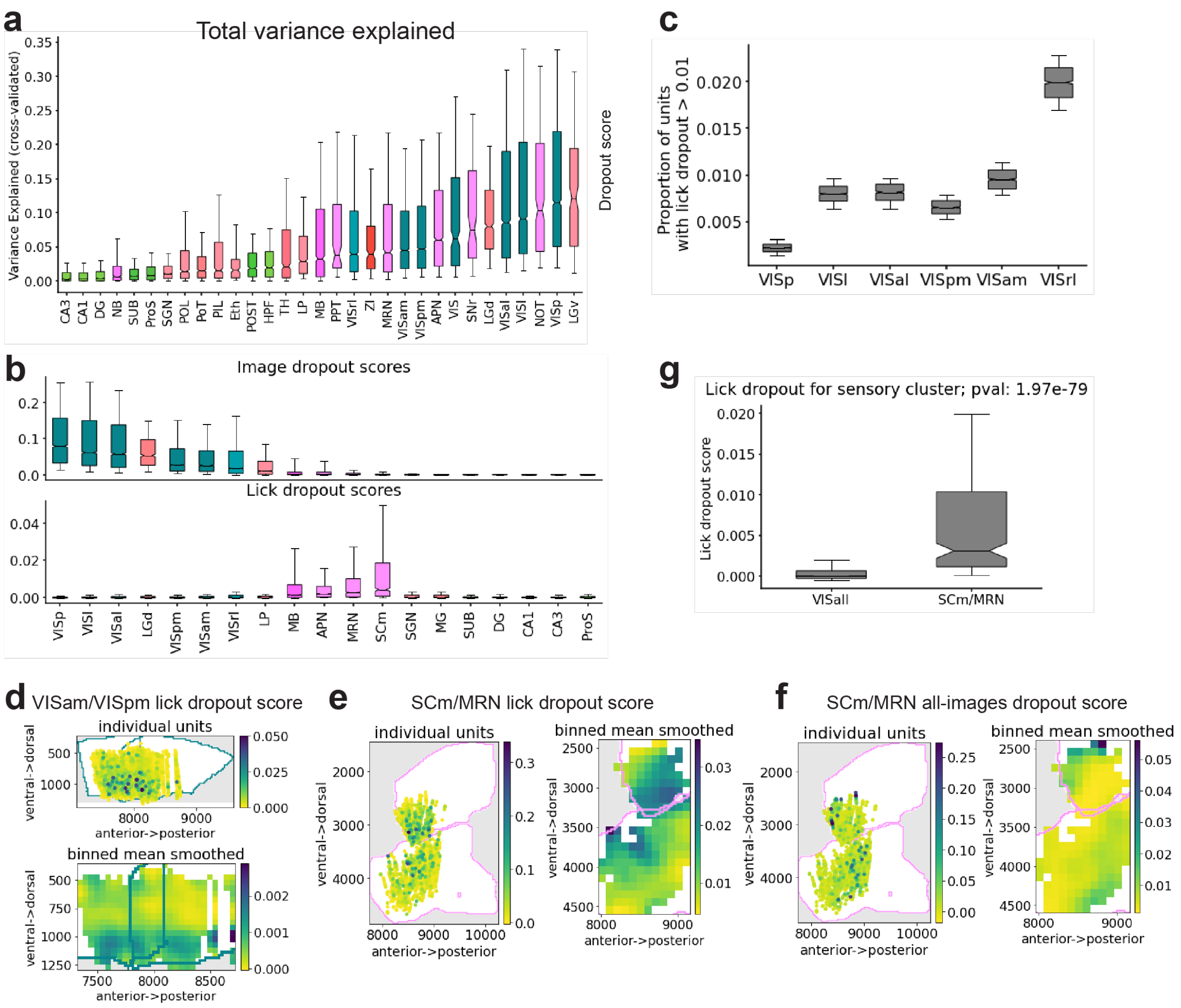


**Figure S3: Further analysis of GLM results. a)** Cross-validated variance explained by GLM fit for units in all recorded areas. **b)** Visual (top) and lick (bottom) dropout scores across recorded areas. **c)** Proportion units with high lick dropout scores across six visual cortical areas. **d)** Distribution of lick dropout scores across cortical depth. Top: Lick dropout scores for each unit recorded in VISpm and VISam collapsed to horizontal plane. Teal contours outline VISam (anterior) and VISpm (posterior). Darker colors indicate larger dropout scores. Bottom: Same data as top, but spatially binned and smoothed. **e)** Same as (d) but for SCm/MRN. Pink contours outline SCm (dorsal) and MRN (ventral). **f)** Same as (e) but for visual dropout scores. Visual dropout scores were calculated by dropping all image kernels. **g)** Lick dropout scores for visual cortical units and SCm/MRN units assigned to sensory clusters. P-value calculated using Wilcoxon rank-sum test. Box plot notches indicate median; box edges indicate 25$^{th}$ and 75$^{th}$ percentiles; whiskers span 10-90$^{th}$ percentiles.

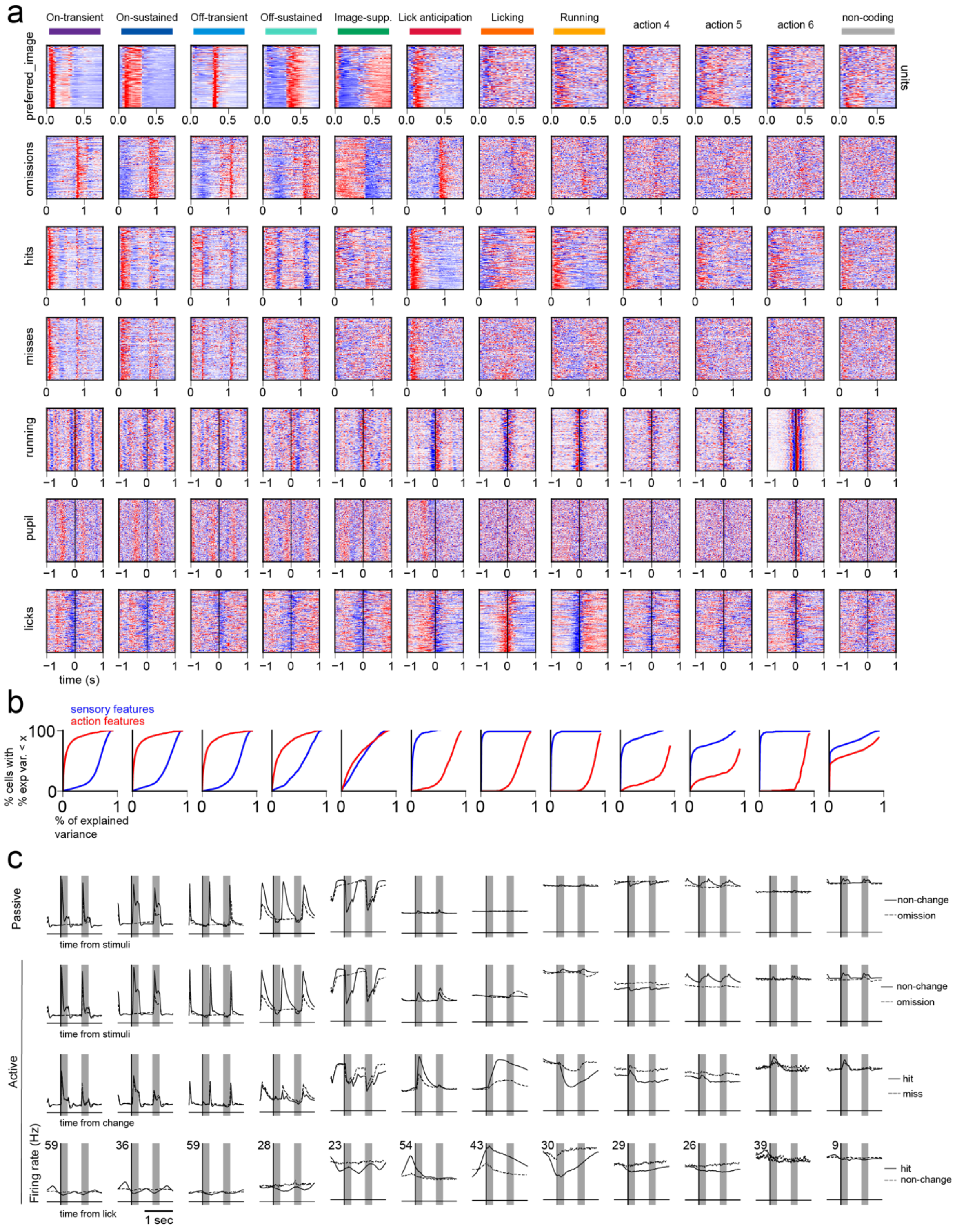


**Figure S4: Kernel fits and population responses for each cluster. a)** Temporal kernels for all units in all clusters. **b)** Cumulative histograms showing what percentage of the explained variance is attributable to sensory (blue) or action (red) features. **c)** Population PSTHs for each cluster aligned to different task events in either the active or passive experimental epoch.

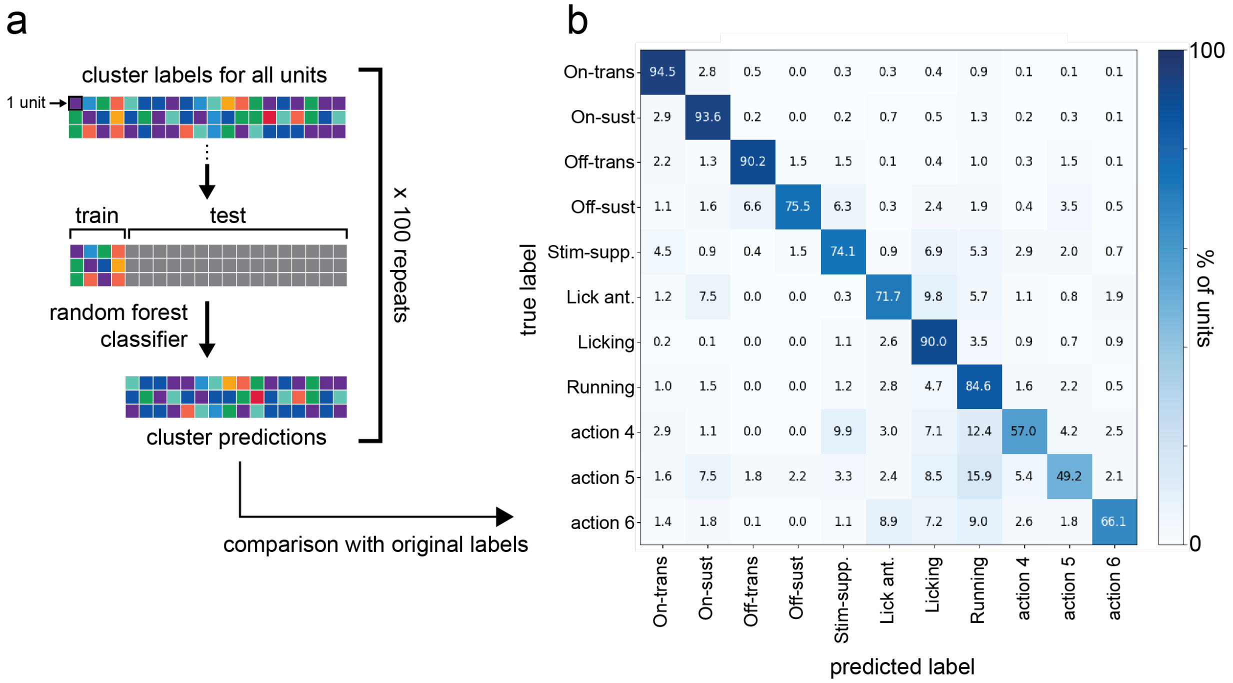


**Figure S5: Cluster validation using bootstrapped classifier.** a) Diagram depicting bootstrapped classifier method. For each iteration (100 total), units were randomly divided into 20/80 train/test splits and a random forest classifier was trained on the cluster labels from the training data to predict the cluster labels of the test set. The input to the classifier was the same feature matrix used in clustering after UMAP dimensionality reduction to three feature dimensions per unit. Predictions were pooled across bootstrap iterations to construct the confusion matrix. b) Confusion matrix indicating the frequency with which units in each cluster were assigned to each cluster by the classifier.

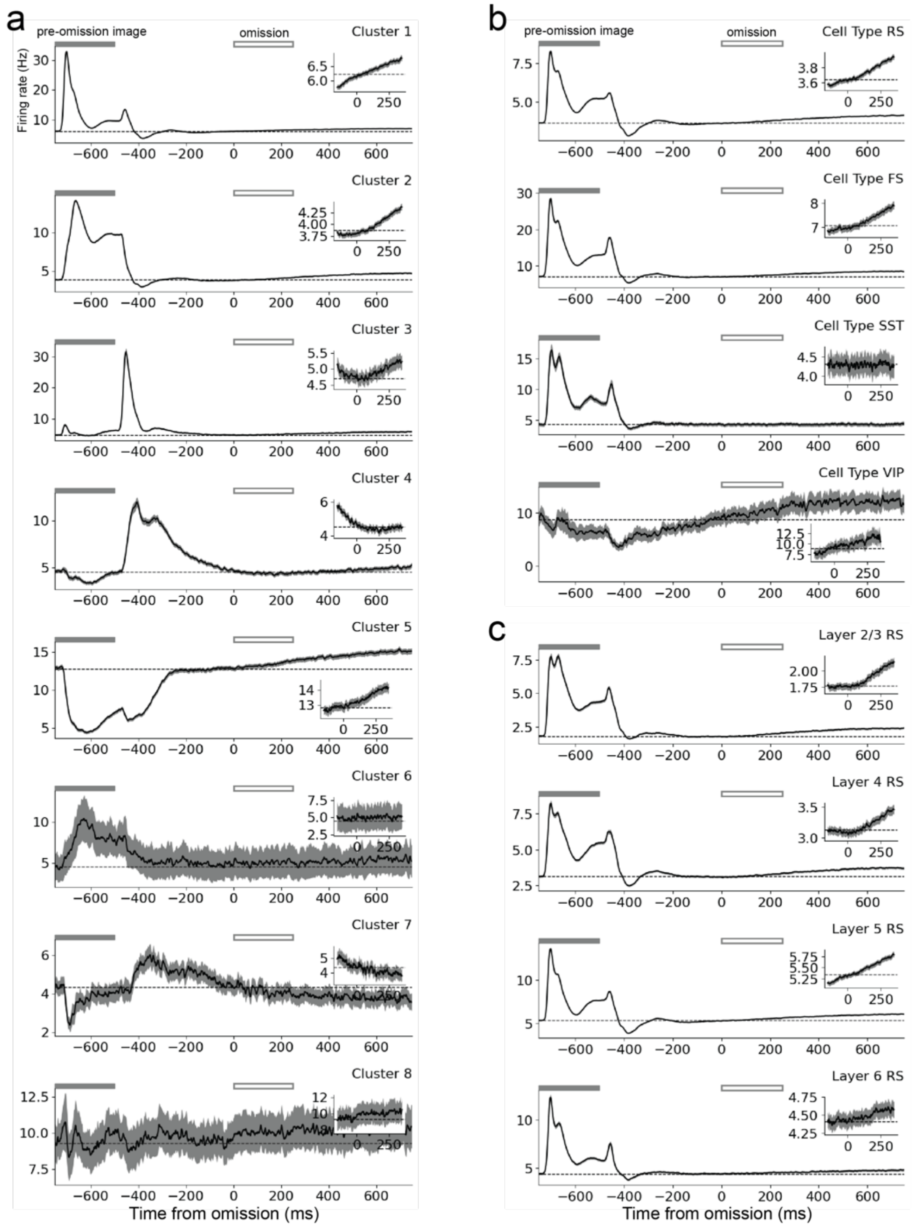

**Figure S6: Response to stimulus omissions in visual cortex. a)** Omission responses for visual cortical units in each of the eight GLM clusters. Filled grey rectangle at top of plot indicates the pre-omission image presentation. Open rectangle indicates when the omitted stimulus would have appeared. Plot inset expands on time of omission. **b)** Same as (a) but partitioning visual cortical units by cell type. **c)** Same as (a) but for RS units partitioned by cortical layer. Shaded regions give SEM.

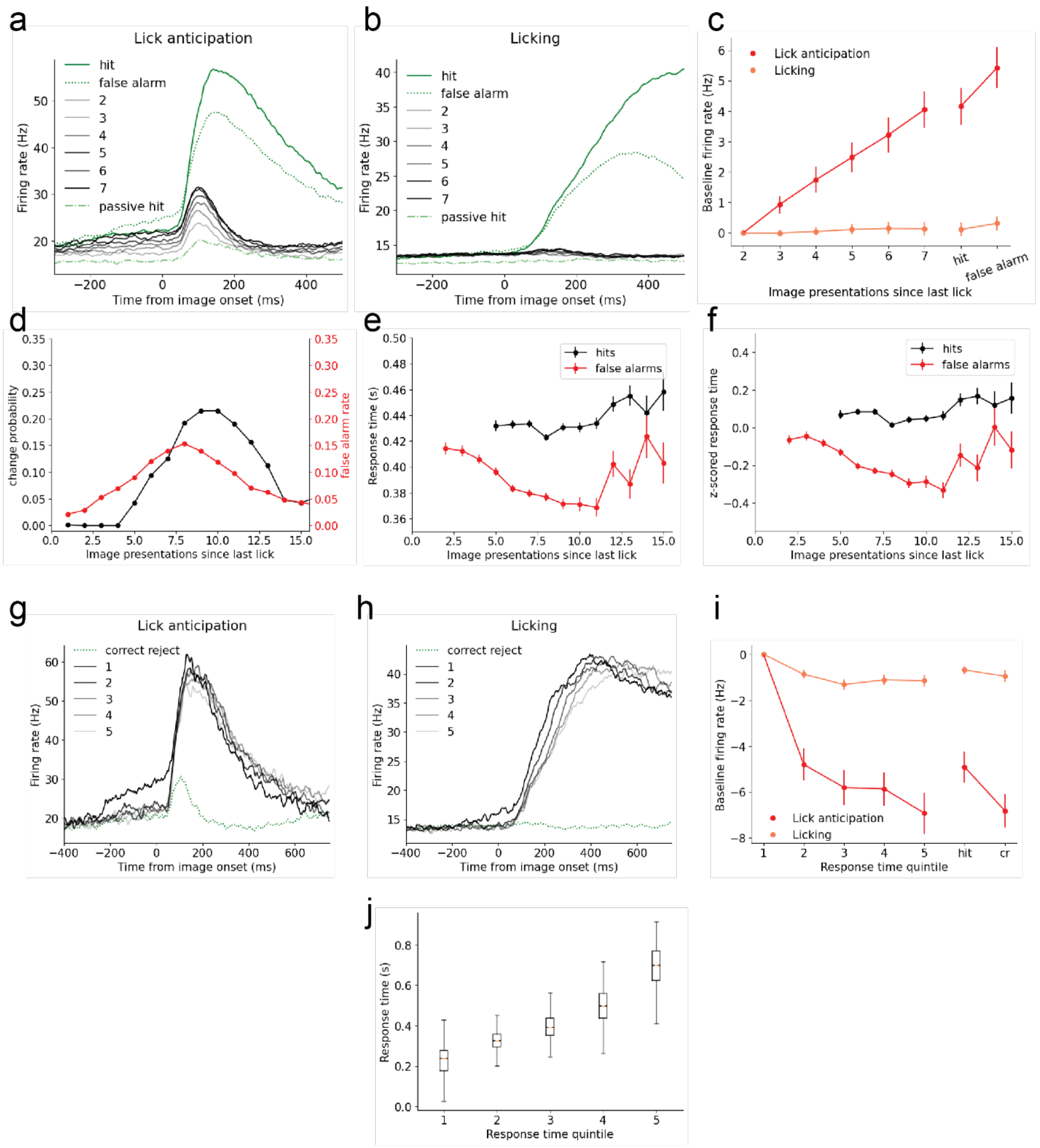


**Figure S7: Baseline modulation of SCm/MRN lick anticipation units. a)** Population response of SCm/MRN units assigned to lick anticipation cluster broken out by the number of stimulus presentations since the last lick response (2-7 in legend) as well as hit and false alarm trials. 'Passive hit' (green, dot-dash) responses refer to change stimuli shown during the passive viewing epoch which correspond to the hit stimuli (green, solid) during the active epoch. **b)** same as (a) but for SCm/MRN units assigned to the licking cluster. **c)** Mean of baseline activity (200 ms before stimulus onset) against presentations since last lick. **d)** Probability of image change (black) and false alarm rate (red) against presentations since last lick.

**e)** Response time as function of presentations since last lick for hit and false alarm trials. **f)** Same as (e) but the response times for each session have been z-scored before averaging across sessions. Error bars are SEM. **g,h,i)** Same as (a,b,c), but now activity is broken out by response time quintile and correct reject trials. Response time quintiles were calculated for each session separately before combining across sessions. Quintile 1 corresponds to trials with fastest response times. **j)** Response times by quintile. Boxes span 25-75$^{th}$ percentile. Orange line indicates median. All active conditions in (a-c) and (g-i) have been matched for mean running speed during the 200 ms prior to stimulus onset.

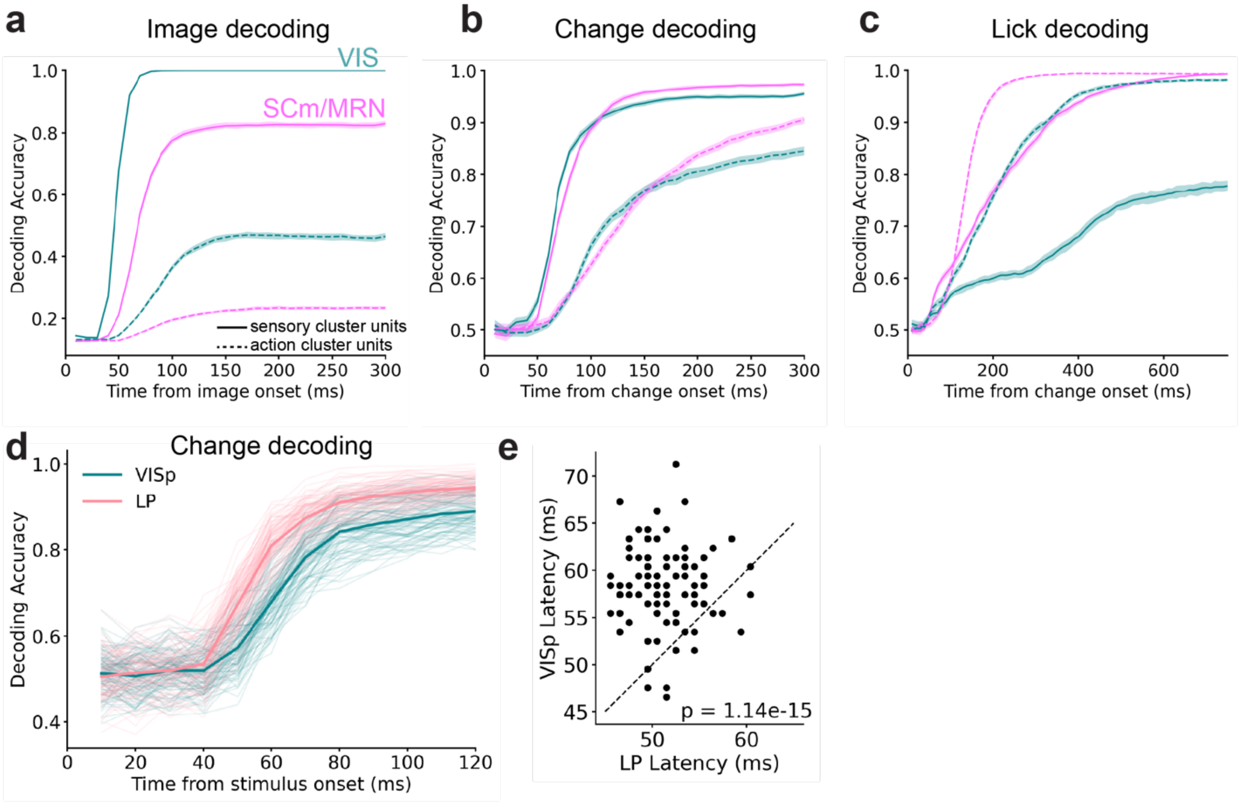


**Figure S8: Comparing decoding time courses across clusters (sensory vs action) and areas (VISp vs LP) a)** Image decoding time courses for VIS (teal) and SCm/MRN (magenta) units from sensory (solid lines) or action (dashed lines) clusters. **b, c)** Same as (c) but for change and lick decoding. Note that action cluster units were too sparse in individual thalamic and cortical visual areas to perform this analysis. **d)** Session-matched change decoding from sensory units in VISp and LP. The number of units from each region were matched for each session before pooling (see Methods). Data from 100 unit-subsampling iterations are shown with thin lines. Thick lines show means across iterations. **e)** Scatter plot of LP and VISp decoding latencies from each iteration (100 total). P-value calculated with Wilcoxon signed-rank test.

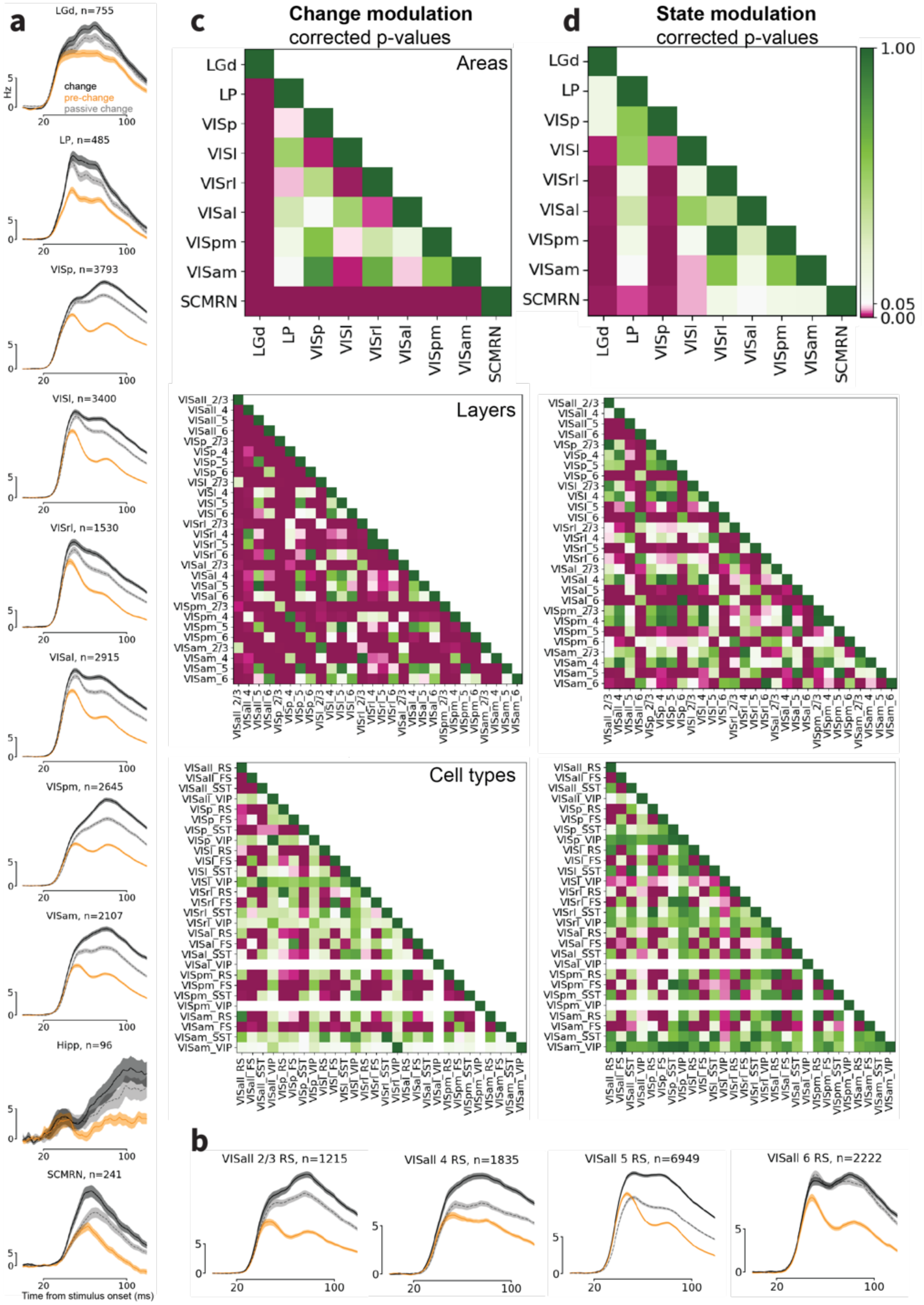


**Figure S9: Change and state modulation across areas. a)** Population responses to change (black) and non-change (gold) stimuli during active behavior. Gray shows change stimuli during passive viewing. **b)**

Same as (a), but plotted for RS cells in each cortical layer. **c)** P-value matrices for change modulation values across areas (top), cortical layers (middle) and cortical cell-types (bottom). All p-values were calculated with Wilcoxon rank-sum test and corrected for multiple comparisons by the Benjamini-Hochberg method. **d)** Same as (c) but for state modulation.

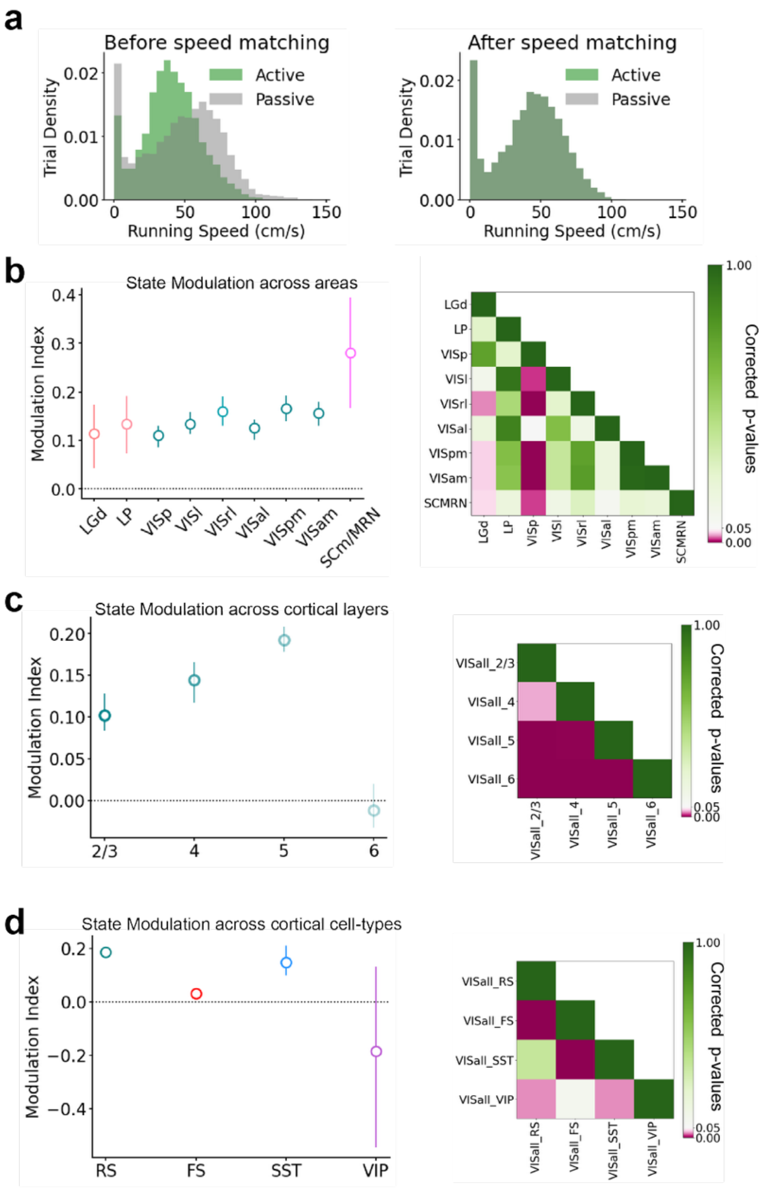


**Figure S10: Matching running speed across active and passive conditions**

**a)** Histogram of running speed during active (green) and passive (gray) epochs before (left) and after (right) speed matching across trials. For each trial, speed was averaged over the 200 ms before change onset. Note that after speed matching, the histograms are perfectly overlapping (dark green color). **b)** Right: State modulation across areas using only speed matched trials. Symbols indicate median and error bars indicate bootstrapped 95% CI of the median. Left: p-value comparison matrix for data on right. **c,d)** Same as (b) but for cortical layers (c) and cell types (d). All p-values were calculated with Wilcoxon rank-sum test and corrected for multiple comparisons by the Benjamini-Hochberg method.

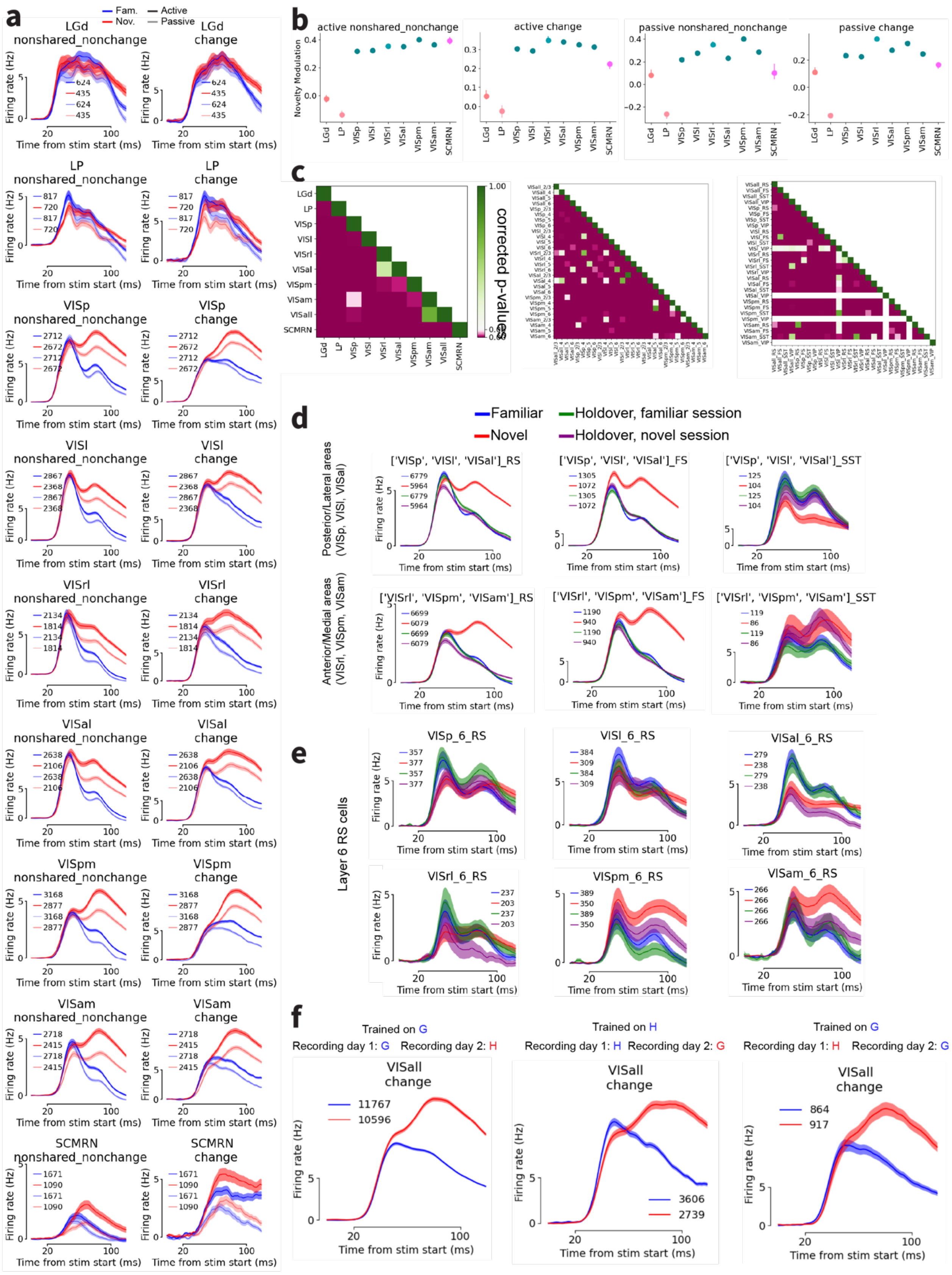

a
Fam.
Nov.
Active
Passive
LGd nonshared_nonchange
LGd change
LP nonshared_nonchange
LP change
VISp nonshared_nonchange
VISp change
VISl nonshared_nonchange
VISl change
VISrl nonshared_nonchange
VISrl change
VISal nonshared_nonchange
VISal change
VISpm nonshared_nonchange
VISpm change
VISam nonshared_nonchange
VISam change
SCMRN nonshared_nonchange
SCMRN change
Firing rate (Hz)
Time from stim start (ms)
b
active nonshared_nonchange
active change
passive nonshared_nonchange
passive change
Novelty Modulation
LGd
LP
VISp
VISl
VISrl
VISal
VISpm
VISam
SCMRN
c
corrected p-values
VISall
d
Familiar
Novel
Holdover, familiar session
Holdover, novel session
Posterior/Lateral areas (VISp, VISl, VISal)
Anterior/Medial areas (VISrl, VISpm, VISam)
['VISp', 'VISl', 'VISal']_RS
['VISp', 'VISl', 'VISal']_FS
['VISp', 'VISl', 'VISal']_SST
['VISrl', 'VISpm', 'VISam']_RS
['VISrl', 'VISpm', 'VISam']_FS
['VISrl', 'VISpm', 'VISam']_SST
e
Layer 6 RS cells
VISp_6_RS
VISl_6_RS
VISal_6_RS
VISrl_6_RS
VISpm_6_RS
VISam_6_RS
f
Trained on G
Recording day 1: G
Recording day 2: H
Trained on H
Recording day 1: H
Recording day 2: G
Trained on G
Recording day 1: H
Recording day 2: G
VISall change
11767
10596
3606
2739
864
917

**Figure S11: Further characterization of novelty modulation. a)** Population responses to non-change stimuli during familiar (blue) and novel (red) sessions for active behavior (dark) and passive viewing (light). For cortical areas, RS population is shown. **b)** Novelty modulation index calculated for change and non-change stimuli during active behavior and passive viewing. Error bars indicate bootstrapped 95% CI for the median. **c)** P-values for novelty modulation indices across areas (left), cortical areas and layers (middle) and cortical areas and cell types (right). All p-values were calculated with Wilcoxon rank-sum test and corrected for multiple comparisons by the Benjamini-Hochberg method. **d)** Population responses for RS (left), FS (middle) and SST (right) units to familiar images (blue), novel images (red), holdover images during the familiar session (green) and holdover images during the novel session (purple). Units are either pooled across VISp, VISl and VISal (top row) or VISrl, VISpm and VISam (bottom row). **e)** Population responses of layer 6 RS units to same stimuli as (d) plotted for each visual cortical region. **f)** Population responses pooled across visual areas for three cohorts experiencing different training and recording trajectories. Note that for all three training/recording trajectories the response to novel stimuli is amplified and delayed relative to familiar stimuli.

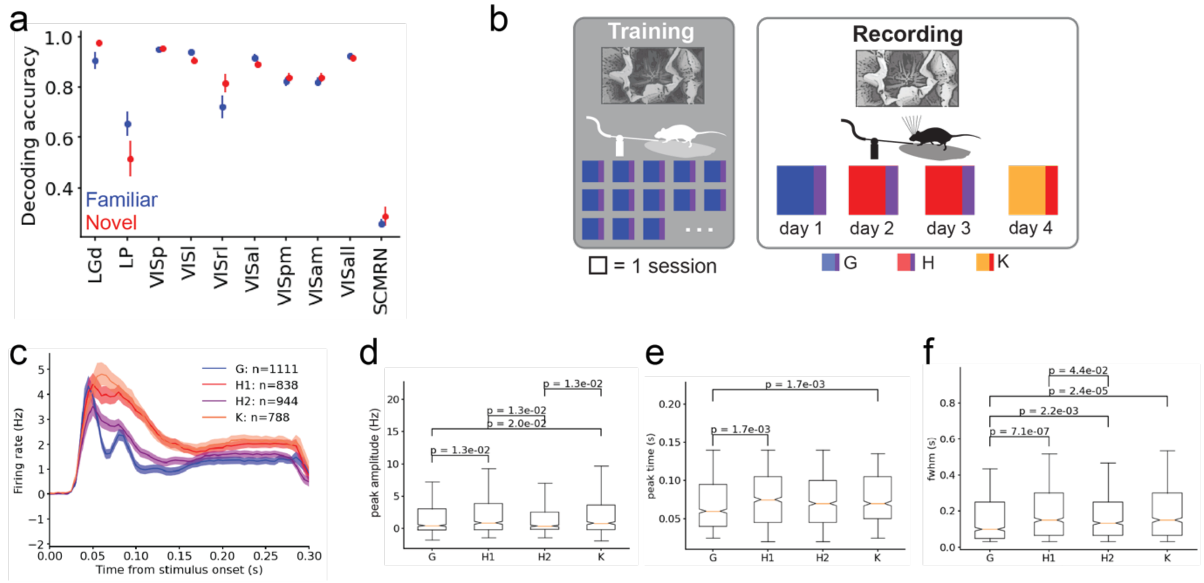


**Figure S12: Time course of image familiarization across sessions. a)** Image decoding accuracy for familiar (blue) and novel (red) images. Error bars indicate SEM across sessions. **b)** Experimental design for four-day experiment: mice were trained on image set G and then recorded for four consecutive days with image sets G, H, H and K respectively. Image set K consisted of six novel images and two holdovers from image set H. **c)** Population responses from visual cortical units to six non-shared images from G (blue), H on recording day 2 (novel, red), H on recording day 3 (repeat, purple) and K (novel, orange). **d)** Peak response amplitude for visual cortical units to images in (c). Notch indicates median. Box indicates 25th and 75$^{th}$ percentiles. Whiskers indicate 10$^{th}$ and 90$^{th}$ percentiles. **e)** Peak response time. **f)** Full width at half max. P-values calculated with Wilcoxon rank-sum test with correction for multiple comparisons.

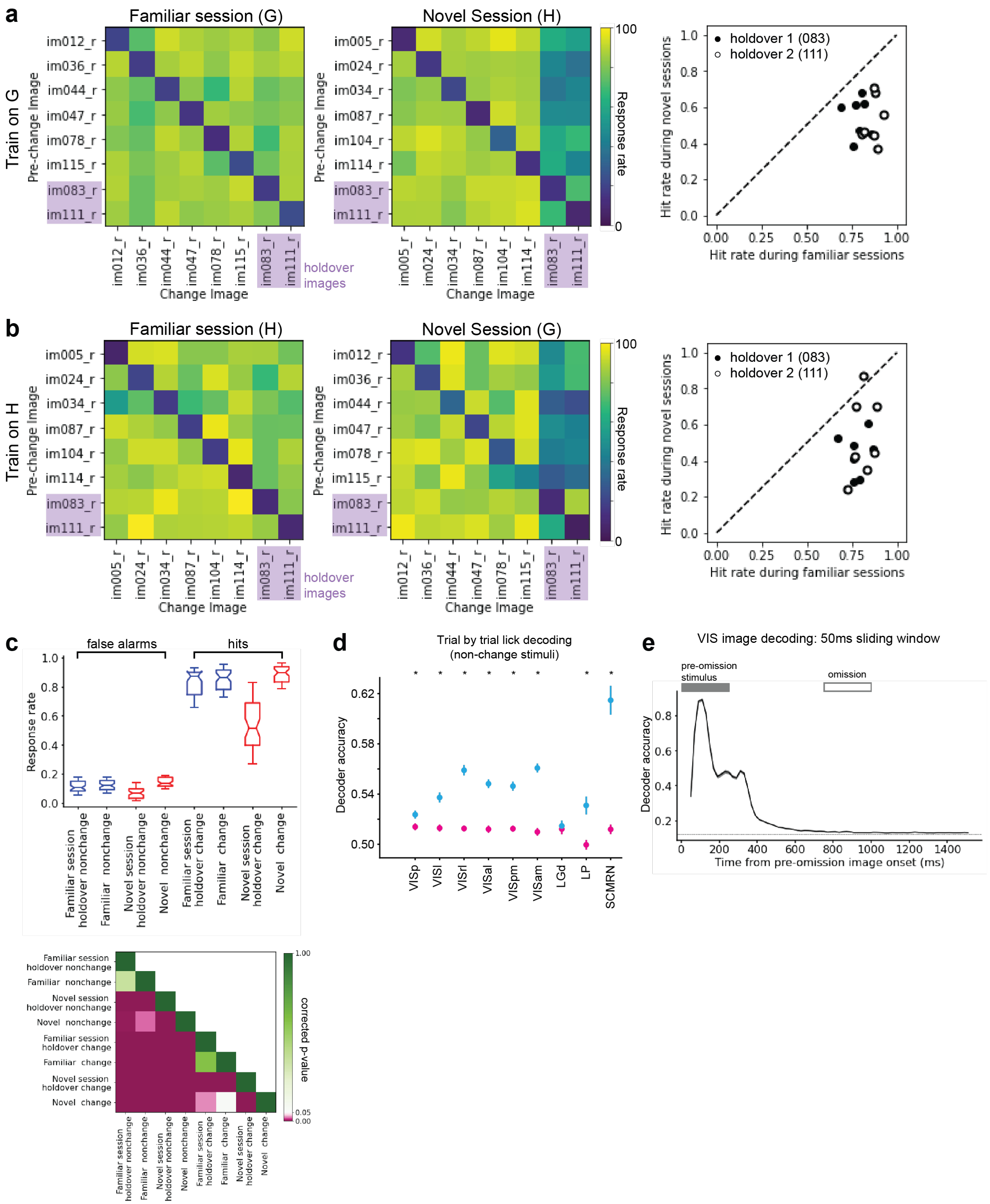

**Figure S13: Visual change detection strategy. a)** Left: Mean response probability for each image transition in image set G for mice trained on G. Holdover images appear in final two rows and columns (purple shading). Middle: Mean response probability for image set H for mice trained on G. Right: Mean response probability for each image transition to holdover images (non-diagonal elements of final two columns) during familiar vs novel session. Image 111 shown in open symbols; image 83 in closed symbols. **b)** Same as (a) but for mice trained on image set H. **c)** Top: Response probabilities during familiar and novel sessions to various stimulus conditions. Bottom: P-value comparisons calculated with Wilcoxon rank-sum test corrected for multiple comparisons. **d)** Lick decoding accuracy during repeat stimuli using the image-comparison decoding strategy or the change decoding strategy. * indicates p <0.05 after correction for multiple comparisons. **e)** Image decoding from visual cortical neurons over time. Decoding was run on activity in a 50 ms sliding window with a new decoder trained for each time step. Pre-omission stimuli were chosen to illustrate how image information decays over the longer inter-stimulus interval. Dotted line indicates chance performance.
